# Functional imaging of hippocampal layers using VASO and BOLD on the Next Generation (NexGen) 7T scanner

**DOI:** 10.1101/2025.08.29.673151

**Authors:** Suvi Häkkinen, Alexander Beckett, Erica Walker, Laurentius Huber, David A. Feinberg

## Abstract

Spatial accuracy and venous biases are a central concern in mesoscale fMRI, with subcortical brain regions facing additional challenges due to lower sensitivity, high physiological noise, and complicated vasculature. Here, we optimized CBV VASO on the NexGen 7T scanner for layer-specific investigations of the human hippocampus. Both VASO and BOLD (from the same acquisition) detected significant hippocampal activation during an established autobiographical memory task. At the macroscale, activation patterns converged, showing pronounced memory task activation in the anterior hippocampus and consistent engagement across the frontoparietal neocortex. At the laminar scale, however, differences in depth-dependent profiles emerged within the subiculum: BOLD exhibited a pronounced bias toward the inner layers, consistent with the known venous drainage pattern in the subfield. Finally, the optimized effective TR of 3.2 s allowed exploratory study of retrieval stages and neocortical functional connectivity, which both supported hippocampal anterior-posterior dissociation using both VASO and BOLD. Thus, hippocampal fMRI allows mapping layer function with high accuracy and can provide deeper insights into a number of neuropsychological phenomena and disorders.

**Key points:**

- Optimized CBV VASO on NexGen 7T for layer-specific hippocampal imaging
- VASO mitigates structural venous bias in subiculum present in BOLD
- Long-axis mapping dissociates retrieval stages and neocortical connectivity patterns

## Introduction

A central pursuit in mesoscale fMRI is to model activation flow in the columns and depths of gray matter (Felleman and Van Essen, 1991). So far, most efforts have focused on neocortical circuits, yet incorporating subcortical areas and circuits would be highly relevant for neuroscientific and clinical applications. Subcortical imaging however faces additional technical challenges related to the longer distance from the receiver coils and high levels of physiological noise. Here we focus on human hippocampus, which has a three-layered organization with gray matter in parts less than 1 mm thick, complex curved shape, and functional specialization along short and long axes (Vos De Wael et al., 2018; Dalton et al., 2019; Zheng et al., 2021; Angeli et al., 2025), with studies emerging also on laminar functional differentiation (Maass et al., 2014; Ahmadi et al., 2024, 2026; Pfaffenrot et al., 2025; Warrington et al., 2025). In addition to the requirement for sufficiently small voxel size and detection sensitivity, hippocampal layer-fMRI faces the challenge of intricate subfield- and layer-dependent vasculature: blood may drain to either inner or outer surface depending on hippocampal region and individual anatomy (Duvernoy et al., 2005). A bias which makes it difficult to reliably localize BOLD activity in neurovascular coupled activation is that blood oxygenation changes are also in the cortical draining veins, creating a confound in differentiating changes in layer activity (Duvernoy et al., 1981; Olman et al., 2007). Therefore, contrast mechanisms that are inherently less sensitive to the large draining veins (Oshio and Feinberg, 1991; Lu et al., 2003; Feinberg et al., 2008) would be very attractive for hippocampal layer-fMRI, but remain challenging in practice.

VAscular Space Occupancy (VASO) contrast (Lu et al., 2003) is able to localize neuronal activity with high spatial specificity based on changes in cerebral blood volume (CBV), primarily the dilation of arterioles, capillaries, and intracortical arteries (Huber et al., 2019), and has been shown to be less sensitive to signal from large draining veins (Jin & Kim, 2008). The VASO contrast and its lesser sensitivity to large draining veins has been validated using animal models and invasive imaging techniques (Huber et al., 2021), and is commonly used in neocortical layer-specific fMRI (Huber et al., 2015, 2017; Beckett et al., 2020). The gain in specificity comes, however, at the cost of a more complex inversion recovery pulse sequence and analysis requiring the difference between two images yielding lower signal-to-noise ratio (SNR) and contrast-to-noise (CNR) of roughly half compared to BOLD at 7T. This is also due to larger venous deoxyhemoglobin dependent signal changes compared to smaller microvascular blood volume changes (Huber et al., 2014, 2015; Beckett et al., 2020). Thus, reaching sufficient detection power with VASO in submillimeter hippocampal imaging is challenging. Further, the pulse sequence and the lower statistical power of the contrast often necessitate the use of relatively long TRs and block designs, limiting the types of neuroscientific inquiries. Recent research conducted on a standard 7T with body gradients (Ahmadi et al., 2025, 2026), simultaneous to this study on the NexGen 7T (Häkkinen et al., 2025), shows promise in applying VASO in hippocampus just as has been shown other low-tSNR brain regions (Rane et al., 2016; Burak et al., 2022; Faes et al., 2023) as well as in rapid short TR experiments (Dresbach et al., 2023). In this work, we will apply VASO to study hippocampal layers on the Next Generation (NexGen) 7T scanner (Feinberg et al., 2023) with technical improvements of several scanner subsystem components specifically adapted for mesoscale imaging at 7T. These include a channel array receiver coil and a head-only asymmetric gradient coil “Impulse” (200 mT/m, 900 T/m/s) for ten times higher performance than the body gradient coil available on standard 7T systems. This advanced hardware on NexGen 7T allows for better optimization of VASO technique for high-resolution imaging in hippocampus, through higher acceleration performance and SNR from the high-channel array, and reduced time of EPI echo trains using the shorter echo spacing allowed by the faster head gradient coil. Thus, the scanner allows the opportunity for VASO sequence optimization.

In the present study, we applied an established autobiographical memory paradigm to assist in the interpretation and validation of VASO contrast results. The recall of autobiographical memories rich in detail and context reliably activates hippocampus in fMRI (Addis et al., 2007; Leelaarporn et al., 2024), providing an effective testbed for comparing VASO and BOLD contrasts and their sensitivity to venous biases which may mislead interpretation at the mesoscale (Pfaffenrot et al., 2025). By comparing VASO and BOLD from the same acquisition, our goals were to: 1) Test the sensitivity of VASO and BOLD in detecting hippocampal activation during an autobiographical memory task. 2) Demonstrate lesser venous bias in VASO compared to BOLD in the inner layers of subfield subiculum, where blood drains through inner layers. 3) Test the possibility of detecting temporal dynamics during memory retrieval, requiring sensitivity to shorter blocks. In addition, using the available neocortical data, we 4) explored hippocampal-neocortical functional connectivity patterns based on VASO and BOLD. In particular, we tested whether both contrasts capture the established relative difference between anterior and posterior hippocampus to large-scale functional networks (Angeli et al., 2025; Borne et al., 2023; Vos De Wael et al., 2018) and if there are layer-specific connectivity differences in the subiculum consistent with venous bias. Together these results will inform on the sensitivity and specificity of VASO and BOLD in the study of hippocampal circuitry.

## Materials and Methods

### Subjects

Ten participants (7 female; age 18–44 years) were scanned for the main analysis, and two additional in vivo scan sessions were used to identify the capabilities and challenges of hippocampal layer-fMRI as well as to implement and validate the imaging protocol. One participant was excluded because they did not finish the session. The study protocol was approved by the office for protection of human subjects, UC Berkeley IRB, and each participant gave written informed consent before MRI data acquisition.

### Autobiographical memory task experiment

The task paradigm was adapted from previous studies on fine-grained hippocampal activation, which have demonstrated stronger activation across hippocampal subfields during *autobiographical memory* (AM) compared to *mental arithmetic* (MA) task (McCormick et al., 2015; Leelaarporn et al., 2024; Pfaffenrot et al., 2025). In the AM task, participants were presented with a cue-word (e.g., coffee) and were instructed to recall an event from their personal past that took place less than three years ago and lasted less than a day. The participants were told to press a button when they had selected a memory and then silently think about the event in detail until the word disappears, as if experiencing it again. Unique cue-words were taken from the Clark and Pavio extended norms (Clark and Paivio, 2004) based on high

Thorndike-Lorge frequency, imageability and concreteness, as originally in (Addis et al., 2007). In MA, participants were presented with a simple mathematical problem (e.g., 15 + 12; additions, subtractions, multiplications, divisions) which they were to solve, then press a button, and start iteratively adding 3 to the result until the instruction disappears. AM and MA tasks were presented in alternating blocks separated by short fixation blocks. Data collection was TR-locked to the beginning of each task or fixation block, and task blocks had duration of 6 TR and rest blocks one TR, leading to 19.3 s task blocks separated by 3.2 s fixation blocks. Functional data was collected in three runs of 11.6 min (total 35 min). One subject was scanned twice on different days (total 70 min) to explore intra-subject test-retest reliability.

Participants briefly practiced the tasks right before the scan. Task compliance during scanning was measured by recording response button presses and verbally confirmed after the scan. Eight out of nine participants indicated their task performance consistently throughout data acquisition (>97% and 88% of trials in the AM and MA task, respectively; **Supplementary Table S1**), but button presses were by mistake not recorded for one participant. Data from all nine participants was included in main analyses comparing AM and MA blocks, but the data from the participant without recorded motor responses was excluded from the task stage comparison.

### Data acquisition

Data were collected on the NexGen 7T scanner equipped with the high performance Impulse head-only gradient coil “Impulse gradient” with maximum amplitude and slew rate (Gmax 200mT/m, SR 900 T/m/s) and a 64-channel receive array coil (Feinberg et al., 2023). CBV weighted functional data were collected using a VASO sequence (Huber et al., 2014) with a skipped-CAIPI 3D EPI readout. On the NexGen 7T scanner, the shorter echo spacings achievable using the Impulse gradient reduced signal dropout in the temporal regions, and importantly allowed readout of the 3D image slab within a single inversion recovery (IR) cycle of blood labeling. Therefore a segmented approach across multiple IR cycles was not required, which reduced the phase artifacts arising from such a segmented acquisition (**Supplementary Figure S1**). The scan parameters were: Matrix Size 206×206×30, field of view (FOV) 175×163, slice thickness 0.84 mm, TE 13.6 ms, In-plane segmentation 3, GRAPPA 1x3_z1_, Bandwidth 1516 Hz, Echo Spacing 0.72 ms, TI1/TI2 939 ms/2097 ms, TR_vol_ 1.158 s, effective TR (nulled/not-nulled) 3.2 s. The FOV was positioned sagittally, covering the right hippocampus and parts of the frontoparietal neocortex (**Supplementary Figure S2**).

Targeting a single hemisphere allowed us to utilize a smaller FOV and a sagittal slice orientation provided in-plane coverage along the long axis of the hippocampus. This approach minimizes phase-encoding steps and echo-train length to achieve the resolution, reduces the required acceleration, and in turn raise temporal SNR, reduces TR, and mitigates susceptibility-induced geometric distortions compared to alternative image orientations (Ahmadi et al., 2026; Huber et al., 2024). The right hippocampus was chosen based on models of functional lateralization, where it is specialized for mapping visuospatial architecture, allocentric coordinate frames, and the mental imagery of scenes (Burgess et al., 2002) as well as has more distributed functional connectivity with large-scale cognitive control and neocortical networks (Robinson et al., 2016). Because our layer-fMRI paradigm relied on intense internal visual imagery, we hypothesized that the right hippocampus may exhibit stronger temporal dynamics when contrasting the phases of active scene construction and steady-state elaboration (Daselaar et al., 2008).

For anatomical reference, a whole brain MP2RAGE scan was also collected (voxel size 0.75 mm isotropic, matrix size 300x300x208, TE 2.82 ms, TR 6000 ms, Partial Fourier 7/8, GRAPPA 3, TI1/TI2 800 ms/2750 ms).

### Hippocampal subfield segmentation and layer extraction

The MP2RAGE was used to estimate hippocampal inner and outer surfaces and segmentation of subfields subiculum, cornu ammonis (CA) 1–4, and dentate gyrus (DG) using HippUnfold (version 1.4.1; DeKraker et al., 2022). The surfaces (with an approximate vertex spacing of 0.5 mm) were then transformed to VASO space to avoid resolution loss due to data interpolation. Registration of VASO and MP2RAGE was initialized by rigid manual alignment. Second, we applied ANTs to compute global affine alignment (mutual information) and a non-linear warp (cross-correlation) for local anatomical refinement (SyN; version 2.5.0; Avants et al., 2008), with 20 iterations at the native voxel scale. The affine and nonlinear transforms were applied to inner and outer hippocampal surfaces to bring them to native functional space using Connectome Workbench (wb_command version 1.5.0, Marcus et al., 2011). All registrations and final inner and outer hippocampal surfaces were manually inspected in functional space. If vascular artifacts (bright inflow in VASO) or B0-inhomogeneities interfered with alignment causing non-anatomical stretching, the registration was iterated with manually drawn cost-function masks excluding these regions and stronger regularization.

For final depth sampling, 21 equi-volume surfaces between inner and outer surface and 5 surfaces extending outside these limits were generated (*wb_command -surface-cortex-layer*). Following conventions of the HippUnfold toolbox and previous hippocampal layer-fMRI literature (Ahmadi et al., 2026; Pfaffenrot et al., 2025), we refer to the layers closest to the stratum radiatum/lacunosum-moleculare (SRLM) boundary as inner and layers closest to stratum oriens/pyramidale boundary as outer.

### Neocortical layer extraction

Sagittal FOV allowed us to collect data from large regions of the frontoparietal neocortex. Available neocortical data was analyzed for task activation and hippocampal-neocortical functional connectivity patterns. Neocortical surfaces were reconstructed from high spatial resolution (0.75 mm isotropic) MP2RAGE scans using the submillimeter recon-all pipeline from FreeSurfer (Dale et al., 1999; Zaretskaya et al., 2018). Analogous to hippocampal surfaces, the white and pial surfaces were carefully registered to VASO with a combination of rigid (BBR; Greve and Fischl, 2009) and nonlinear transforms (ANTs SyN), and manually quality-controlled for neocortical alignment. 25 equivolume surfaces were generated representing different cortical depths.

### Functional data preprocessing and quality assessment

Nulled and not-nulled series were motion-corrected to mean volume of the first run (AFNI 3dVolreg). Noise reduction with distribution corrected PCA (NORDIC PCA; version 4/22/2021; Moeller et al., 2021; Vizioli et al., 2021) was applied to nulled and not-nulled images separately, on magnitude only (Knudsen et al., 2025). VASO contrast data was generated using dynamic division (Huber et al., 2014), which involves temporal upsampling to better align the series of images with and without blood nulling, and was also applied to BOLD data. This provided VASO and BOLD time-series data acquired in the same acquisition, which we then analyzed separately using the same pipeline. Because BOLD and VASO signal changes are opposite in sign, the VASO signal was inverted for easier comparison. To ensure magnetization reached a steady state, the first 8 upsampled volumes of each contrast were discarded from every run.

To characterize within-study data quality, temporal SNR (tSNR) was calculated across all conditions by dividing the temporal mean by temporal standard deviation. Region of interest (ROI) values were extracted in native voxel space for the full hippocampus, which was defined using the HippUnfold *multihist7* atlas to encompass the subiculum, CA1–CA4, and DG. Reproducibility across runs of the same participant was tested using intraclass correlation coefficient (ICC; Shrout and Fleiss, 1979) using pingouin package (Vallat, 2018) and two-way mixed-effects modeling, treating subjects as random effects and runs as fixed effects.

### Functional activation analysis

Activation analyses were performed in native native voxel space using general linear models (GLM) as implemented in Nilearn (Abraham et al., 2014) and managed via Nipype workflows (Gorgolewski et al., 2011). Analyses used two designs. The first design matrix had two regressors of interest (AM, MA), and a contrast was defined to contrast the two tasks. The second, three task regressor design, further split AM condition into separate construction and elaboration stages based on the button press during the trial, and a linear contrast compared these two memory task stages. For the second design, only trials with button press were modeled, and the one participant (S3) without recorded button presses was not analyzed. The average time of construction and elaboration stages were 3.2 and 16.2 s, respectively (**Supplementary Table S1**). Both models used a high-pass filter of 0.011 Hz. To confirm that results are not explained by physiological noise, we repeated analyses including the six head motion parameters, outliers (> 0.3 mm mean framewise displacement; Power et al., 2012), and physiological correlates from the aCompCor (Behzadi et al., 2007) variant described in (Pfaffenrot et al., 2025). This variant includes signals from a white-matter ROI (a 4 voxel cube manually placed in adjacent white matter; components chosen to explain 50% variance) and a high residual ROI (> 3 SD of all residuals from the GLM fit; 5 components) putatively sensitive to respiratory and cardiac cycles. The aCompCor regressors were orthogonalized with respect to motion regressors. For the AM > MA contrast, the first-level GLM included 26–62 regressors in the primary analyses and 35–89 regressors in the aCompCor control analyses, evaluated against 428 upsampled volumes (corresponding to 214 original timepoints) per run.

For group level assessment, individual hippocampal surfaces were registered to a canonical surface space using HippoMaps (0.1.0, DeKraker et al., 2025) and beta parameters were sampled from volume to each hippocampal surface (*wb_command -volume-to-surface-mapping*) using trilinear interpolation.

Statistical comparisons were conducted with increasing granularity using ROI-based analyses, surface analysis, and continuous layer comparisons. First, an ROI-based approach was taken to compare activation patterns defined by three depth bins and three longitudinal segments (Dalton et al., 2019; Poppenk et al., 2013; Hackert et al., 2002; Leelaarporn et al., 2024). Depth bins were defined by binning the five innermost, middle and outermost layers. Longitudinal segments were defined by equal 33% geometric thirds of the longitudinal axis of the unfolded coordinate system. Activation patterns were compared using a ROI-based 3-way 3 × 3 × 2 repeated-measures ANOVA with Segment (Anterior, Middle, Posterior), Layer (Inner, Middle, Outer), and Modality (VASO, BOLD) as within-subject factors. Significant effects were assessed using post hoc one-sample t-tests with FDR-correction within modality (Benjamini & Hochberg, 1995). Statistics were computed using statsmodels (Seabold et al., 2010).

Finer-grained topological patterns along the hippocampal surface were assessed using surface-based analysis. To account for anatomical variability and increase sensitivity to cluster statistics, surface overlays were smoothed using a 5 mm Full Width at Half Maximum (FWHM) Gaussian kernel (*wb_command -metric-smoothing*). Task activation patterns were analyzed using non-parametric permutation testing via the Permutation Analysis of Linear Models (PALM) toolbox (Winkler et al., 2014). To map memory task engagement, an intercept-only one-sample t-test was computed. To evaluate spatial and laminar magnitude dissociations across depth profiles, within-subject paired t-tests were executed. Statistical inference was conducted on the canonical hippocampal surface manifold (0.5 mm grid density) using Threshold-Free Cluster Enhancement (TFCE; Smith & Nichols, 2009) and standard standard 2D surface parameters (H = 2, E = 1, C = 26). For the one-sample t-tests, the null distribution was constructed using tail-specific sign-flipping. For the paired two-sample t-tests, permutations were restricted within-subject to respect the repeated-measures design. Family-wise error rate (FWER) correction was achieved via 10,000 permutations using one-tailed inference (p_FWE_ < 0.05).

Finally, continuous layer activation profiles were evaluated separately for each subfield. The specificity of VASO and BOLD in detecting layer activation was assessed based on z-scores at different depths of gray matter. Group-average profiles were generated by averaging vertex values per subfield and layer. The shapes of VASO and BOLD depth profiles (z-scores) were then explicitly compared at group using the Generalized Additive Model (GAM) framework as implemented by the AFNI PTA tool (Chen et al., 2021) utilizing the mgcv R package (Wood and Scheipl, 2020). This framework treats depth as a Multilevel Smoothing Spline (MSS), allowing the evaluation of continuous morphological trajectories across the hippocampal depths without discrete depth binning. Specifically, we modeled the smooth main effect of layer depth and the interaction between layer and condition (Contrast) as fixed effects, while incorporating inter-subject variability as a random intercept term via s(Participant, bs=“re”). Smoothing parameters were optimized via Restricted Maximum Likelihood (REML). To capture non-linear profile geometry while preventing overfitting, the maximum basis functions for the splines were constrained to k = 4. Differences between the BOLD and VASO trajectories across the continuous laminar gradient were explicitly evaluated by testing the significance of the Contrast × Layer interaction spline. CA4 and DG were excluded from all layer-specific analyses since the neuronal laminar organization of these subfields is less clear. Results were FDR-corrected across the four investigated subfields.

### Functional connectivity analysis

Connectivity analysis assessed connectivity patterns between areas of the hippocampus and available neocortex. We expected functional connectivity based on VASO and BOLD to show similarity to the spatial patterns of task-evoked activation. In particular, we expected that the available parts of the frontoparietal attention/control and default mode networks would show stronger activation during math and memory tasks, respectively, delineating functionally specialized areas. Based on previous literature, the anterior and posterior hippocampus should also have clear relative differences in their connectivity to the frontoparietal and default mode networks. Finally, we expected that depth-specific connectivity patterns might differ between VASO and BOLD, particularly in subiculum which has venous drainage via inner layers.

Connectivity was assessed following the “background connectivity” approach (Fair et al., 2007; Gratton et al., 2018). Data was preprocessed by regressing out task effects (blocks) and nuisance covariates (aCompCor, six motion parameters, motion outliers of > 0.3 mm FD), and high-pass filtering (0.01 Hz), after which outlier volumes were removed from the remaining timeseries. After this preprocessing in the upsampled space, the concatenated runs retained a minimum 1,017 total residual volumes. To account for temporal interpolation, this scales conservatively to the equivalent of 500 independent residual timepoints, providing approximately 27 minutes of clean data. Time series data was sampled to all hippocampal layers and averaged in groups of five layers to represent the inner, middle and outer depths. Seed time series was defined by averaging vertices based on subfield and longitudinal segment. As the subiculum is a major output region of the hippocampus with neocortical circuits arising primarily from the pyramidal cell layer, we sampled the subiculum middle layer for this network assessment. Connectivity was measured as Pearson correlation between hippocampal seed time-series and brain voxels. For each subject, volumetric results were sampled to inner or outer neocortical surfaces, smoothed 3 mm FWHM on surface to account for individual variability (*mri_surf2surf*), and normalized (fsaverage) using spherical surface registration (Fischl et al., 1999). Group-level patterns were assessed using non-parametric permutation testing as for activations.

## Results

### Hippocampal VASO and BOLD activation to autobiographical memory task

To assess the capability of VASO and BOLD in detecting hippocampal activation, we measured activation during an established autobiographical memory task paradigm. Following NORDIC denoising, full hippocampal temporal signal-to-noise ratio (tSNR) values demonstrated stability across functional runs. Absolute VASO tSNR averaged 20.40 (16.65–25.72) across the cohort with excellent run-to-run replicability (ICC(C,1) = 0.96, p < 0.0001). Similarly, the corresponding BOLD timecourses yielded a mean tSNR of 30.50 (26.82–37.10) and remained highly reproducible across runs (ICC(C,1) = 0.84, p < 0.0001). For tSNR values resolved per subfield, see **Supplementary Figure S3**.

Contrasting autobiographical memory and mental arithmetic conditions revealed that, at macroscale, VASO and BOLD patterns converged in hippocampus as well as neocortex (**Figure 1**). At group level, both imaging modalities successfully captured hippocampal engagement, with robust signal increases observed in native voxels for both VASO (t_8_ = 4.272, p < 0.01) and BOLD (t_8_ = 6.182, p < 0.001).

**Figure 1.**
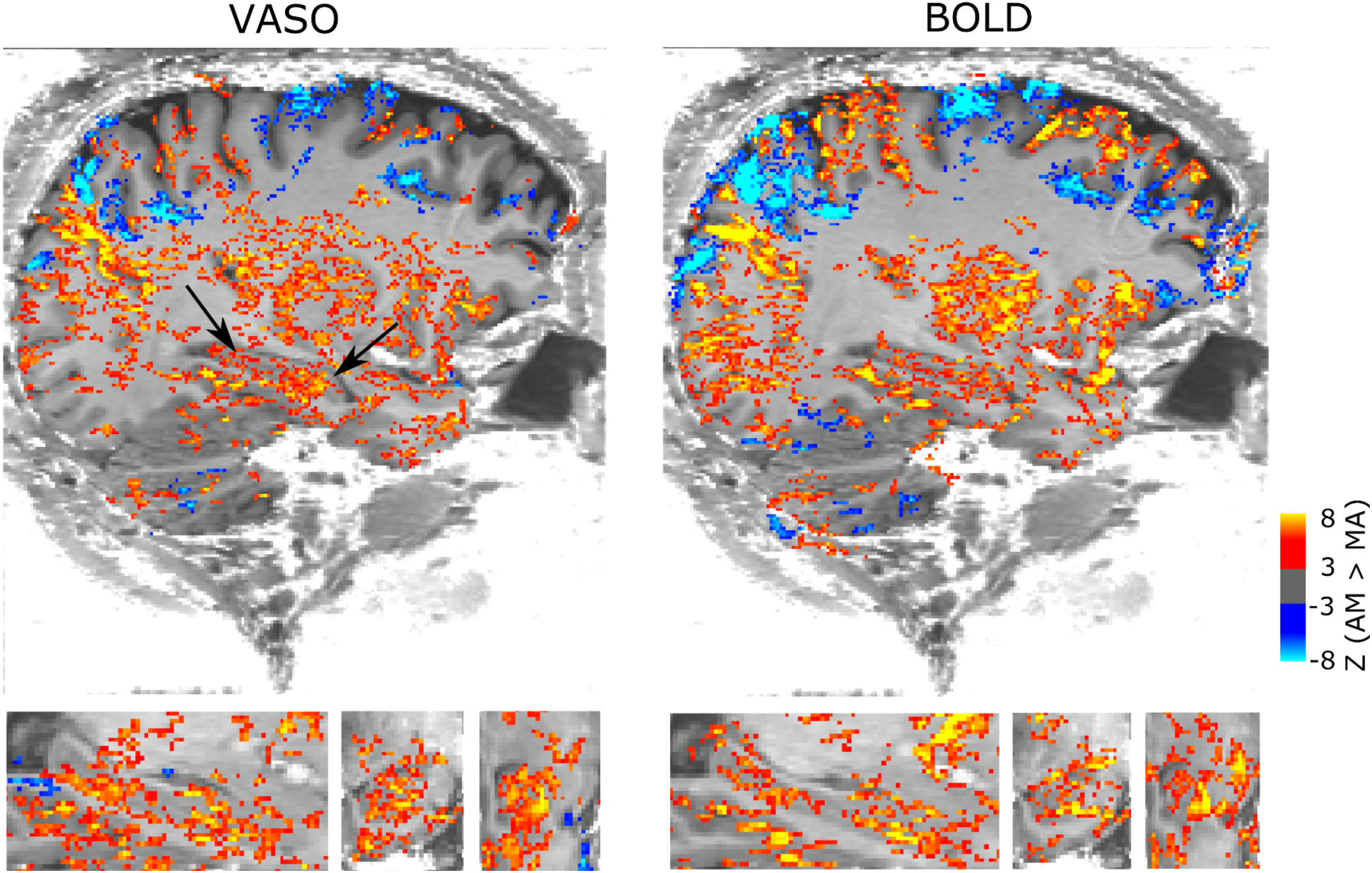
Consistent macroscale activation for VASO and BOLD during memory task (AM > MA). This analysis was based on two sessions (70 minutes) of data. For this macroscale visualization only, VASO and BOLD were denoised with NORDIC factor error levels of 1.5 and 1.0, respectively, to highlight the noisier VASO pattern. Both were thresholded at Z > 3 and clusters of 100 voxels (AFNI *3dClusterize*).

The hippocampus was then segmented by subfield, layer and longitudinal axis (**Figure 2A–B**). VASO and BOLD activation patterns were compared using an ROI analysis using a 3-way 3 × 2 × 2 repeated-measures ANOVA evaluated Segment, Layer, and Modality as within-subject factors (**Figure 2C**; see **Supplementary Table S2** for complete statistics). The omnibus model revealed significant two-way interactions for both Segment × Modality (p = .030) and Layer × Modality (p = .006), alongside significant main effects for all three individual factors (all p < .029). The 3-way interaction did not reach statistical significance (p = .945). To unpack the significant two-way interactions, follow-up paired t-tests were conducted (Holm-Bonferroni corrected within modality). For the Segment × Modality interaction, pairwise comparisons revealed that the long-axis profile variation was driven by differences in BOLD between the Anterior and Posterior segments (t_8_ = 3.191, p_Holm_ = .038), whereas no longitudinal segment pair differences reached statistical significance for VASO. For the Layer × Modality interaction, BOLD demonstrated significant differences across all progressive laminar transitions—including Inner vs. Middle, Middle vs. Outer, and Inner vs. Outer (all p_Holm_ < .034). Conversely, laminar transitions within the VASO profile did not survive multiple comparison corrections. In sum, while both modalities suggested activation peaking in the inner layers, these laminar differences and the strong anterior emphasis were more pronounced in BOLD.

**Figure 2.**
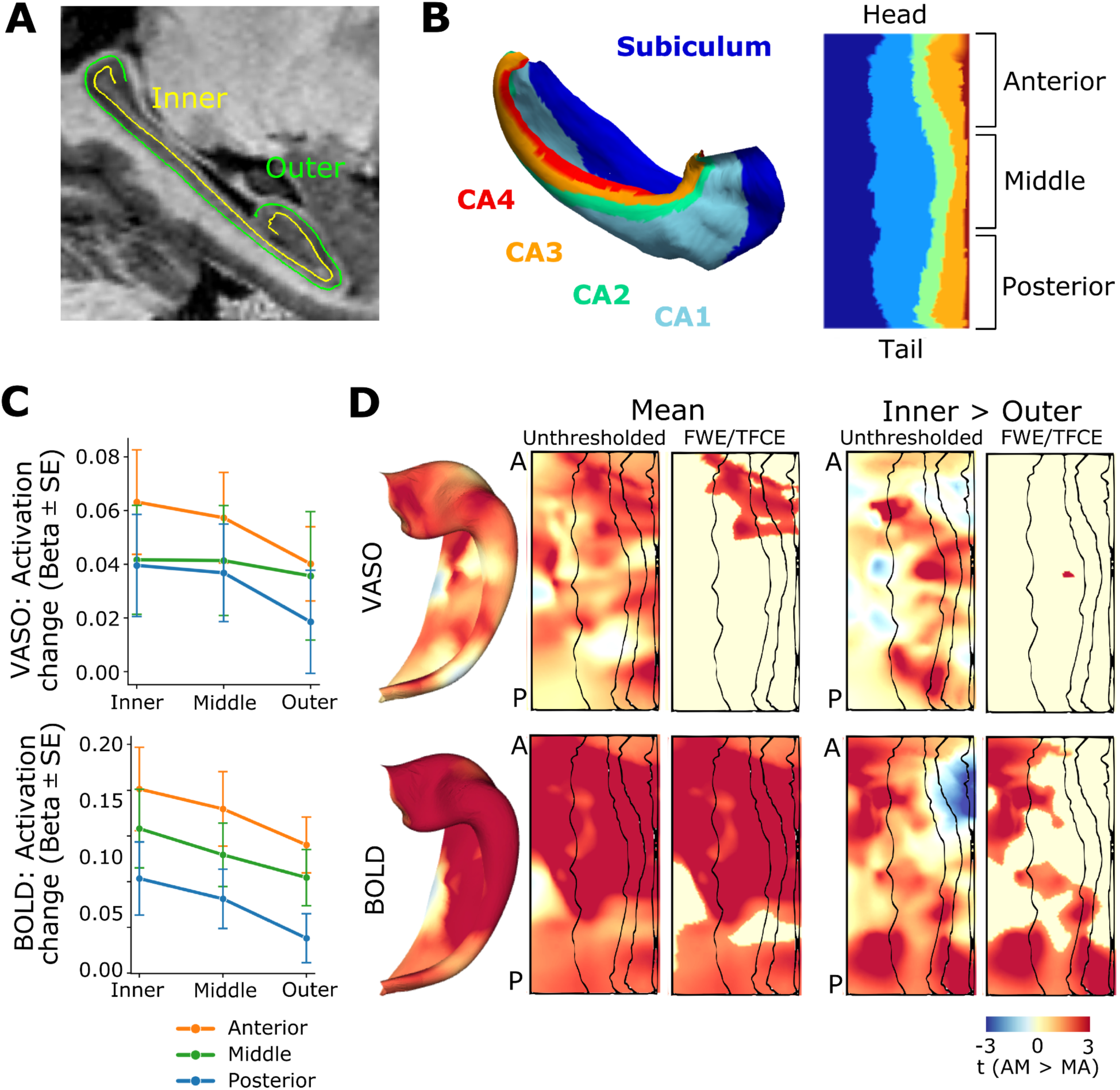
Hippocampal memory task activation (AM > MA) based on VASO and BOLD from the same acquisition. (**A**) Hippocampal inner and outer layers overlaid on sagittal MP2RAGE. (**B**) Hippocampal subfield segmentation on folded and unfolded hippocampal surfaces, as well as the division to anterior, middle and posterior thirds along the longitudinal axis of the unfolded hippocampus. (**C**) Memory task activation (AM > MA) tracked across longitudinal segment (anterior, middle, posterior) and depth (inner, middle, outer) bins. 3-way repeated measures ANOVA (Layer, Segment, Modality) confirmed significant interactions of Layer × Modality and Segment × Modality. (**D**) Group activation patterns visualized on unfolded hippocampal surfaces (5 mm surface smoothing). Within each modality, unthresholded t-statistic maps are displayed side-by-side with family-wise error rate thresholded maps. Panels display general activation topology averaged across gray matter depth alongside maps contrasting inner and outer depths. Both VASO and BOLD showed increased activation in the anterior hippocampus during the memory task. Clusters with stronger activation in inner than outer activation were more extensive in BOLD compared to VASO.

Consistent with these results, vertexwise analysis revealed clusters of anterior hippocampal activation (AM > MA) using both VASO and BOLD (**Figure 3D**). BOLD activation showed extensive clusters with differing inner and outer depth activation, which were not present in VASO results. This divergence between modalities was qualitatively prominent within the subiculum, where VASO did not show similar depth-dependent difference even at uncorrected thresholds. For individual-level maps, see **Supplementary Figure S5**.

**Figure 3.**
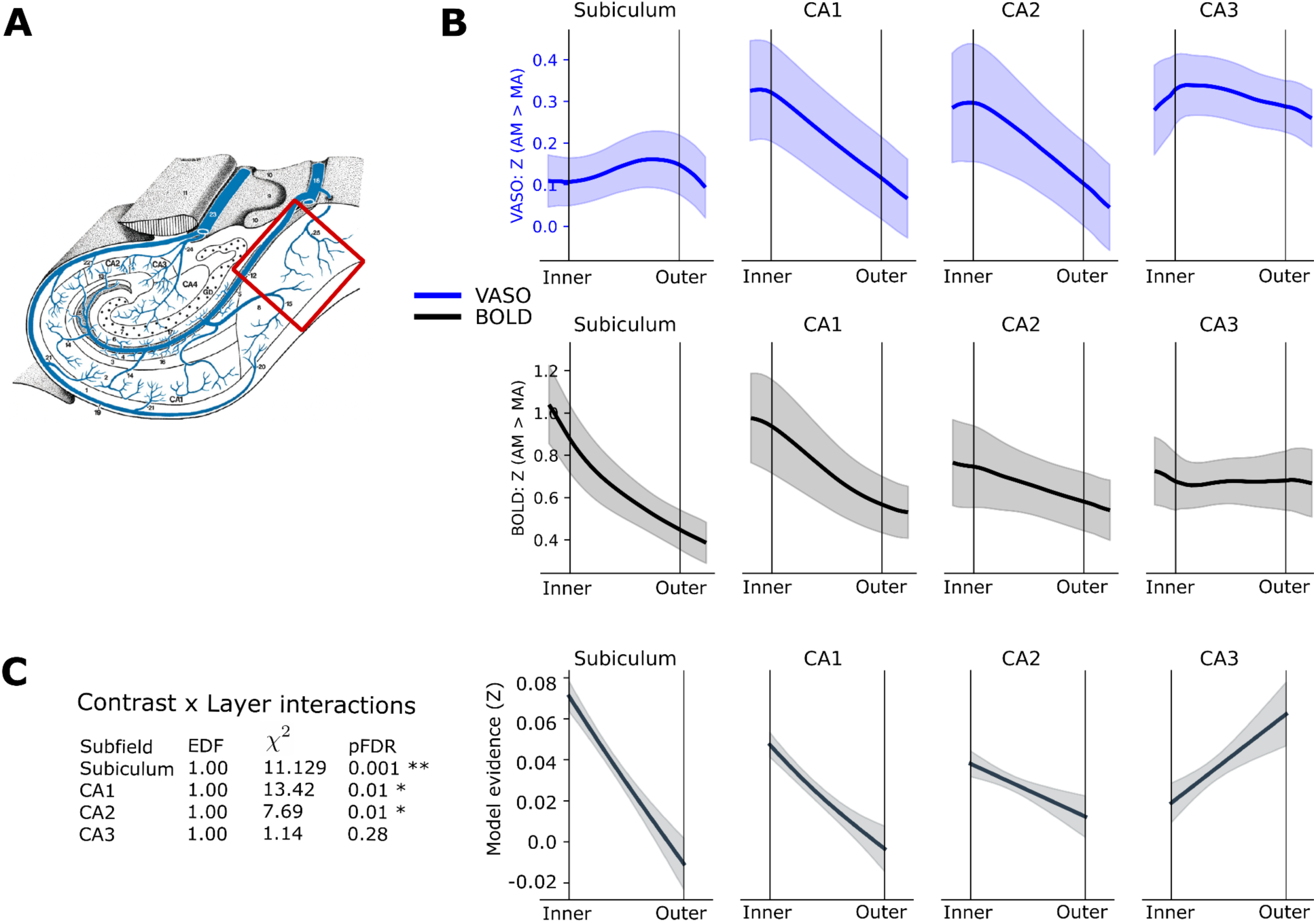
Layer-specific memory task activation (AM > MA) based on VASO and BOLD. (**A**) Schematic of hippocampal venous drainage patterns (modified from Duvernoy et al., 2005), with the inner layers’ venous bias in subiculum highlighted. (**B**) Mean activation depth profiles (z-scores) plotted continuously for VASO (blue) and BOLD (black) across anatomical subfields. Note the stronger activation z-scores in the inner layers of the subiculum in the BOLD but not VASO results, consistent with a venous bias. In CA1, stronger inner layer activation is suggested by both contrast mechanisms. Thick lines denote the group mean and shaded areas represent standard error of the mean (SEM). (**C**) ROI-based PTA analysis, comparing the shapes of VASO and BOLD layer profiles using multilevel smoothing splines, revealed significant differences in subiculum, CA1 and CA2. Model evidence curves support higher BOLD than VASO activation in the inner layers of the subiculum. The shaded ribbon represents pointwise standard error from the population-level model prediction.

Finally, continuous depth profiles of activation (AM > MA) were extracted separately for each subfield. In line with the known venous drainage pattern in the subiculum towards the inner surface (**Figure 3A**; Duvernoy et al., 2005), group-average activation profiles revealed stronger BOLD than VASO activation during the memory task in its inner layers (**B**). This subicular VASO–BOLD difference was seen quite systematically also at individual participant level, with an activation increase in BOLD that was absent or reversed in VASO (**Supplementary Figure S4**). The shape discrepancies were confirmed at group level by PTA analysis (**Figure 3C**), which statistically evaluated whether the two contrasts exhibit a significant difference in profile shape after accounting for baseline amplitude differences. The descending model evidence curves localize the gray-matter depth of greatest divergence sharply to the inner layers for the subiculum, indicating that the statistical confidence for a shape mismatch is concentrated at the inner depths.

The PTA analysis indicated systematic differences between the VASO and BOLD layer profiles also in CA1 and CA2. In both subfields, the shape difference was characterized by a steeper decline from inner to outer layers in VASO. In CA3 the difference was not significant.

The relationships of VASO and BOLD activation profile shapes were quite robust to preprocessing choices. In particular, re-running all analyses with physiological noise correlates estimated via aCompCor revealed highly consistent results (**Supplementary Figure S6**). In the ROI-based repeated-measures 3 × 3 × 2 ANOVA, identical interactions remained significant and post-hoc testing revealed a matching pattern of effects, with the sole exception that the BOLD middle-to-posterior difference reached significance. Unfolded surface analysis similarly revealed highly consistent clusters: VASO activation clusters emerged as slightly more extensive within the anterior hippocampus, whereas BOLD activation and inner-to-outer laminar differences were less extensive in the posterior third. PTA analysis confirmed significant differences between VASO and BOLD depth profiles in subiculum, CA1 and CA2 also using aCompCor, with highly similar slopes across the same subfields as our primary pipeline.

In the current study, we applied NORDIC denoising at the default factor noise level of 1.0. Based on testing in our first subjects, NORDIC greatly facilitated the detection of hippocampal VASO activation profiles by increasing z-scores without fundamentally changing layer profile shape (**Supplementary Figure S7**). Higher factor noise levels began to affect depth response shapes; for example, reducing effect size of the expected inner layer peak in CA1 (Ahmadi et al., 2026; Pfaffenrot et al., 2025). This may result from signal components not being separable from the thermal noise in high noise datasets (Kay, 2022; Olesen et al., 2023; Faes et al., 2024).

Finally, analysis of the test-retest data available on one subject revealed a similar VASO–BOLD activation difference in the inner layers of the subiculum and similarity in CA1 in both 35 min sessions (**Supplementary Figure S8**). Pooling these sessions together increased the resulting z-scores without altering the underlying depth profile shape.

### Hippocampal activation during memory elaboration vs. retrieval stages

We explored activation differences between the construction and subsequent elaboration stage of autobiographical memory recall, known to have partially distinct neuronal correlates. The methodological challenge with this comparison is the speed of these dynamics, as reconstruction duration is similar to our TR of 3.26 s (“AM response time” in **Supplementary Table S1**).

An ROI analysis using a 3-way 3 × 3 × 2 repeated-measures ANOVA evaluated Segment, Layer, and Modality as within-subject factors (**Figure 4A**; see **Supplementary Table S3** for complete statistics). The omnibus model revealed a highly significant main effect of Segment (F_2,14_ = 7.236, p = .007), whereas the main effects of Layer and Modality did not reach statistical significance. Crucially, no two-way or three-way interactions were significant, indicating that the activation pattern across the long-axis did not differ across layers or imaging modalities. In follow-up pairwise comparisons, both imaging modalities demonstrated stronger elaboration activation in Posterior compared to Anterior segments (p_Holm_ < .011) and VASO additionally captured significantly stronger activation in Posterior compared to Middle segment (p_Holm_ = .037). Control analysis with aCompCor reiterated these results except for reaching significance on Segment × Modality interaction, and in post-hoc testing a significant difference between Middle and Posterior segments in VASO but not BOLD. In sum, both VASO and BOLD consistently demonstrated pronounced activation during elaboration compared to construction in the posterior third of the hippocampus, with an uniform profile across layers.

**Figure 4.**
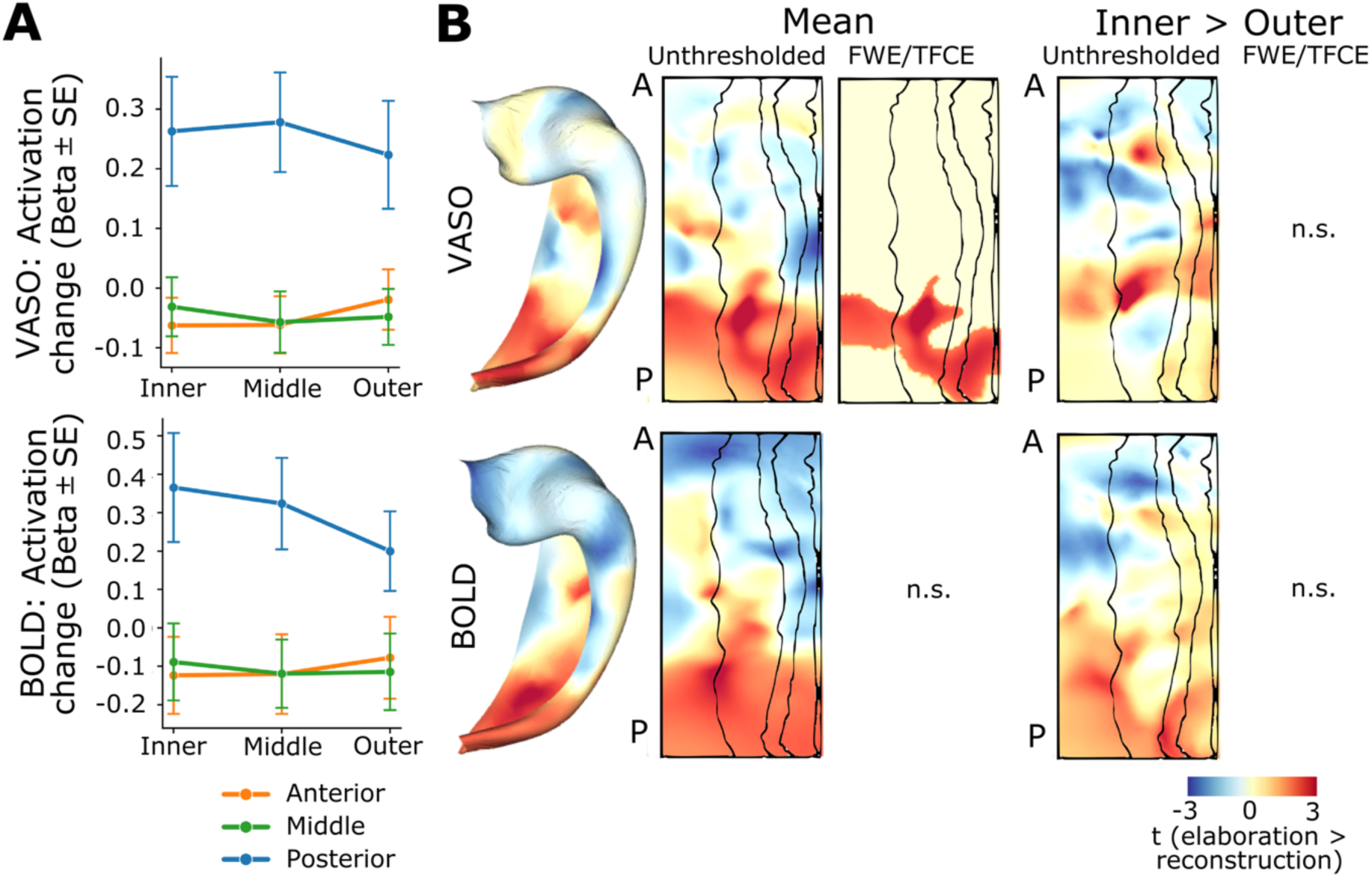
Exploratory analysis of memory retrieval stage dependent activation (elaboration > construction) based on VASO and BOLD from the same acquisition. (**A**) Memory task activation (AM > MA) tracked across longitudinal segment (anterior, middle, posterior) and depth (inner, middle, outer) bins. 3-way repeated measures ANOVA (Layer, Segment, Modality) revealed the main effect of Segment, associated with stronger activation in the posterior hippocampus during elaboration compared to construction. (**B**) Group average activation patterns visualized on unfolded surfaces (5 mm surface smoothing), with VASO indicating significant clusters of stronger activation during elaboration in posterior hippocampus. (**C**) Comparison of inner and outer layer activation was not significant when controlled for multiple comparisons in either contrast.

Surface-based analyses comparing activation to memory task stages indicated significantly stronger activation for elaboration than construction for posterior hippocampus using VASO (**Figure 4B**; for individual maps, see **Supplementary Figure S9**). BOLD activation showed clusters in the posterior hippocampus at uncorrected thresholds (p < 0.05). Layer-dependent differences were not significant. Control analysis with aCompCor regressors only affected the extent of the significant clusters.

### Neocortical VASO and BOLD activation to autobiographical memory task

The right neocortical surface demonstrated fine-grained task-activation patterns during the memory task (AM > MA; **Figure 5A**), with further functional anatomical and functional characterization via probabilistic atlases. Memory-related elaboration activation peaked in the middle frontal gyrus (red; HCP_MMP1 areas 8Ad and 46; Glasser et al., 2016), which probabilistically maps to the default mode network (specifically DefaultC; Yeo et al., 2011), as well as visual and motor cortices. Conversely, relatively stronger math-related activation was observed in posterior parietal regions, spanning the medial and lateral intraparietal areas (blue; e.g., HCP_MMP1 areas MIP, LIPd) mapping onto the dorsal attention and executive control networks (DorsAttnA and ContB).

**Figure 5.**
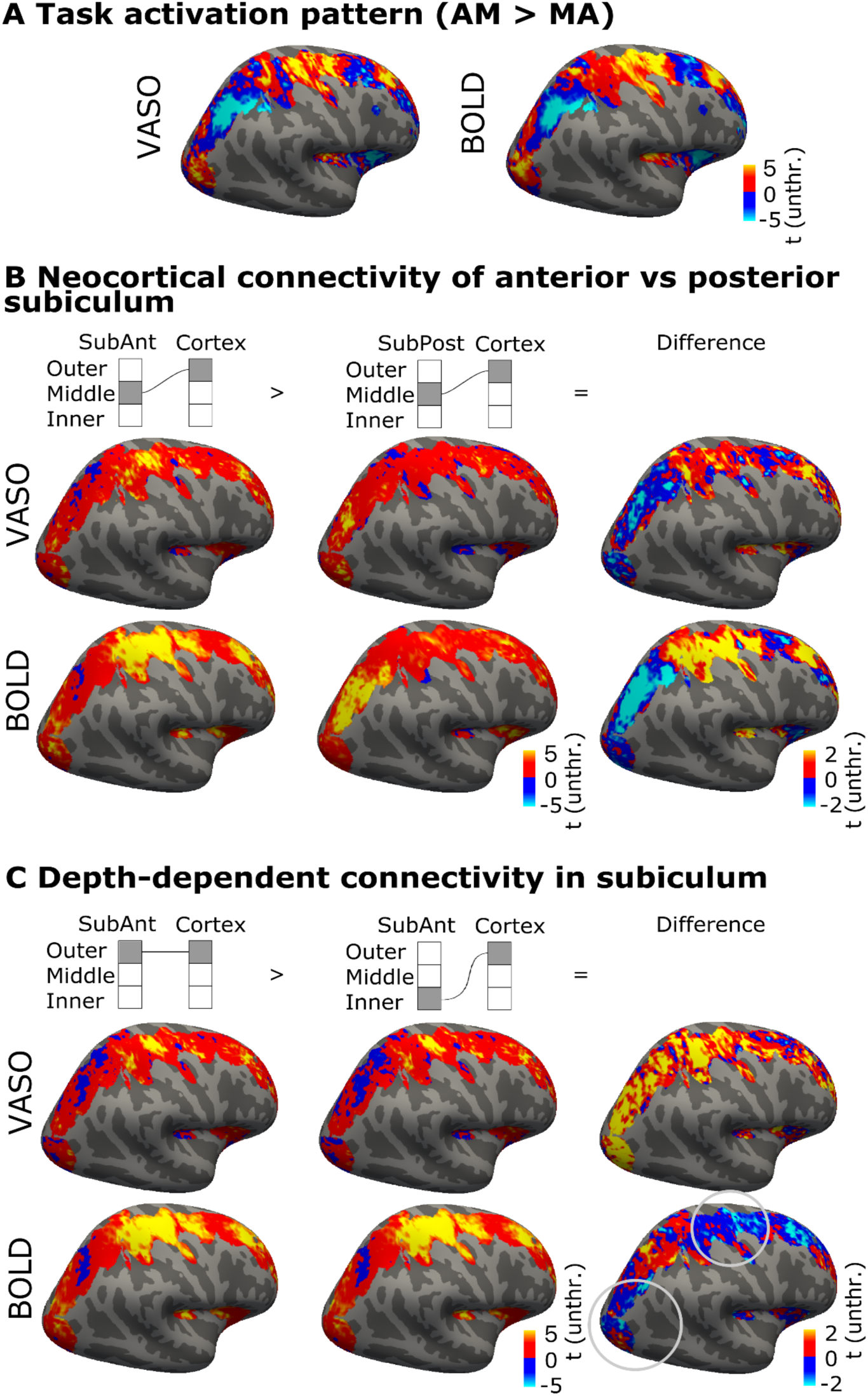
Neocortical activation and hippocampal functional connectivity patterns based on VASO and BOLD. (**A**) Converging activation patterns (AM > MA) indicated functionally distinct areas. (**B**) Functional connectivity patterns associated with anterior and posterior subiculum, a major output region of the hippocampus, showed relative anterior–posterior differences. (**C**) Connectivity seeded by inner and outer depths of anterior subiculum, a region with a known outer-to-inner venous bias. VASO and BOLD layer connectivity differed particularly in areas associated with memory task activation (circled), where BOLD showed stronger connectivity associated with the inner layers of subiculum. In (**A**–**C**), statistics are shown only for the right hemisphere vertices imaged in all nine participants. All statistics are shown without thresholding or correction for multiple comparisons.

### Hippocampal neocortical connectivity

Connectivity analysis focused on the relative difference between the anterior and posterior hippocampus, and the presence of venous bias in the subiculum connectivity. As a comparison point, the functional delineations were expected to show some resemblance to the patterns of task activation (**Figure 5A**). Both VASO and BOLD based connectivity indicated that the posterior subiculum has relatively stronger connectivity to areas more strongly activated for the math than memory task (blue regions in **Figure 5B**), whereas the anterior subiculum had stronger connectivity to memory-related activation (red regions in **Figure 5A**). This spatial pattern was clearer in BOLD than VASO, likely explained by the lower SNR in VASO.

Depth-dependent connectivity profiles within the subiculum indicated a divergence between modalities (**Figure 5C**; for other subfields see **Supplementary Figure S10**). The largest discrepancies between BOLD and VASO emerged within neocortical regions engaged by the memory task (red in **Figure 5A**). Within several of these regions, BOLD connectivity suggested a relatively stronger inner relative to the outer subiculum connectivity, whereas VASO demonstrated the reverse profile (stronger connectivity from outer layers). Thus, the difference might be driven by the venous signal reflected in BOLD based connectivity, misplacing signal fluctuations from outer to inner layers of subiculum leading to overestimation of inner-layer activity.

## Discussion

This study demonstrates that CBV VASO is an effective tool for investigating hippocampal circuitry, and demonstrates the advantages in using advanced imaging hardware for the acquisition of such data, in the context of an established autobiographical memory task. Concurrently acquired VASO and BOLD differed in the suggested layer-resolution localization of activation and connectivity, consistent with a dominant venous signal in BOLD obscuring depth-dependent distinctions in activity. While several task activation effects were observed quite similarly using both contrasts, others were obscured by the venous biases in BOLD, producing clear mislocation (subiculum) and possible artifactual amplification (CA1) of task effects of interest. VASO is therefore a strong alternative for studies on circuit mechanisms where cortical depth-specificity is critical.

### VASO and BOLD differences in activation are consistent with venous bias in hippocampus

VASO and BOLD results converged well at the macroanatomical level, indicating activation in the hippocampus during autobiographical memory retrieval. Task-evoked activation was overall stronger in the anterior compared to posterior hippocampus (**Figure 2**), but also changed dynamically with posterior parts showing stronger activation during elaboration compared to construction stage of the task (**Figure 4**). As expected, however, VASO and BOLD results differed at layer resolution.

The dual contrast fMRI approach used here with VASO and BOLD extracted from the same acquisition allowed us to identify which BOLD layer profiles can be interpreted with respect to neuronal laminar circuitry and which profiles need to be considered as vascular biases. In particular, we planned to use subiculum as a test case, as previous literature is clear that the blood in subiculum is drained through veins along the inner surface (Duvernoy et al., 2005), with respective expected artifactual fMRI signal misplacement (Haast et al., 2024; Pfaffenrot et al., 2025). The activation results in the subiculum clearly show a VASO–BOLD mismatch with more activation in the inner layers using BOLD than VASO, implying less bias in VASO (**Figure 3B–C**). Thus, a BOLD based analysis might lead to the interpretation that input-receiving outer layers of the subiculum are less active, and the inner output layers more active, than they truly are.

Contrasting with the clearcut difference between VASO and BOLD results in the subiculum, results were quite different in CA1, which also has a prominent (outer to inner) vascular bias. In CA1, both VASO and BOLD contrasts indicated stronger activation in inner than outer layers (AM > MA; **Figure 3B**), suggesting that there is a true task-related activation difference between depths of CA1. This finding aligns with the previous layer-BOLD study (Pfaffenrot et al., 2025), which compared responses to the autobiographical task and hypercapnic challenge, and found them both associated with greater CA1 inner layer activation but with distinct laminar response shapes. They concluded that while venous bias is not the sole driver of this inner layer CA1 response, results in the area should be interpreted with caution. While VASO is less sensitive to large-vessel effects than BOLD, it is not a ground-truth measure of neuronal activity and remains sensitive to microvascular architecture. Because venous drainage and expected activation are co-directional in CA1, this convergence across modalities cannot rule out residual microvascular or large-vessel biases shared by both contrasts. Nonetheless, this prominent inner-layer weighting is highly consistent with known anatomical wiring: the major associative inputs to CA1 arise from the trisynaptic pathway (DG → CA3 → CA1) via the Schaffer collaterals, which selectively terminate within the deeper laminae (stratum radiatum and pyramidale; Amaral and Witter, 1989). This pathway is heavily implicated in pattern completion and the retrieval of highly detailed episodic narratives (Rolls, 2013), which directly matches the cognitive demands of our autobiographical memory task.

Our profile shape analysis also indicated systematic differences between the VASO and BOLD layer profiles in area CA2, localized to a shape mismatch in the inner layers (**Figure 3C**). While this may also reflect drainage via inner layers (Duvernoy et al., 2005), the subfield is commonly assumed to have drainage both via inner and outer layers, and any population level biases should be confirmed in a larger sample.

### Neocortical activity and hippocampal connectivity

Hippocampus and neocortex are known to function in concert during memory retrieval. Our FOV captured task-relevant neocortical areas and we were able to explore the hippocampal-neocortical interplay in the right hemisphere. Autobiographical memory was associated with stronger activation in areas likely overlapping with the putative default mode or parietal memory networks (**Figure 5A**, red), strongly heavily implicated in autobiographical retrieval (DiNicola et al., 2020; Kwon et al., 2025), as well as within visual and motor cortices, which may reflect sensory reactivation during vivid recall (Hofstetter et al., 2012; Gilmore et al., 2021). The mental arithmetic task, in turn, elicited stronger VASO and BOLD activation in areas such as the intraparietal sulcus (blue), which is specifically tied to numerical processing and calculation (Dehaene et al., 2004).

Beyond the subfield-specific differences, several of our findings reiterate functional differentiation along the anterior-posterior long axis. First, both contrasts indicated pronounced memory task activation (AM > MA) in the anterior hippocampus (**Figure 2C–D**). The explorative comparison of elaboration and construction stages of retrieval suggested further nuance: activation during elaboration was relatively stronger in the posterior than anterior hippocampus (**Figure 4**). The posterior hippocampal emphasis to elaboration is consistent with previous studies contrasting these task stages (Audrain and McAndrews, 2022; Pfaffenrot et al., 2025) as well as with the theory that posterior hippocampus supports the retrieval of fine-grained details and anterior hippocampus coarser, gist-like memory features (Poppenk et al., 2013; Brunec et al., 2018; Audrain and McAndrews, 2022).

Finally, functional connectivity analyses replicated the relative differences in the connectivity of the anterior and posterior hippocampus to neocortical networks. In this context, the anterior emphasis in the hippocampus for autobiographical memory retrieval is consistent with previous literature suggesting stronger anterior than posterior hippocampal connectivity to the default mode network (Vos De Wael et al., 2018; Tang et al., 2020; Zheng et al., 2021; Borne et al., 2023; Angeli et al., 2025). These studies have reported that the posterior hippocampus, in turn, exhibits relatively stronger connectivity to parietal memory, visual, and sensorimotor networks. Intriguingly, this intrinsic network architecture estimated from task-regressed data closely mirrors our task-activation findings. Specifically, the observation that the elaboration stage engaged the posterior hippocampus more robustly than the construction stage (**Figure 4A**) aligns with the expected involvement of the parietal memory network, which is sensitive to familiarity and goal-oriented cognition. Further study of these hippocampal–neocortical connections might help better characterize these memory-related networks.

To summarize, neocortical (macroscale) task activation based on VASO and BOLD converged (AM > MA; **Figures 1** and **2**). Spatially similar neocortical patterns also emerged in exploratory analyses of hippocampal functional connectivity (**Figure 5**, **Supplementary Figure S7**). Importantly, however, depth profiles of functional connectivity based on VASO and BOLD exhibited qualitative differences within the subiculum, suggesting a relative shift in BOLD toward pronounced neocortical coupling with the inner layers. Given that this descriptive trend aligns with the expected direction of potential BOLD signal displacement in the subiculum, it serves as a valuable cautionary example of how venous confounds in BOLD may distort spatial profiles also in functional connectivity analyses.

### Impact of NexGen 7T scanner on mesoscale functional imaging

These methodological advances have exciting implications for noninvasive, in-vivo mapping of the microcircuits to and from the hippocampus in humans. Optimization of VASO imaging in the hippocampus required us to leverage the advantages of the high signal at ultra-high field and the unique hardware of the NexGen 7T scanner (Feinberg et al., 2023) to achieve high-resolution CBV-weighted imaging with sufficiently short TR to resolve the different subphases of autobiographical memory retrieval, and obtain sufficient SNR to overcome the reduced sensitivity of VASO when compared to BOLD. On standard 7T scanners with slower body gradients, one or more of parameters including TR would need to be compromised for VASO imaging in this subcortical brain region. For example, in comparison to a previous study optimizing VASO sequence for hippocampal imaging on clinical 7T MRI scanners (Ahmadi et al., 2026), the sequence optimized for NexGen 7T achieved similar voxel size and coverage at nearly half the TR per volume pair (3.26 vs 5.66 s) and without needing to acquire data across multiple inversion cycles and the accompanying artifacts.

Of note, the present study is a proof-of-concept study highlighting a sequence optimized for imaging one hippocampus. Our modest sample size (N = 9) is typical for high-field layer-fMRI sequence benchmarking but can limit wider generalizability. The focused FOV allowed us to balance demanding spatial and temporal resolution envelopes. Imaging both hippocampi would fundamentally alter this optimization problem, necessitating an increase in either voxel size or volume TR. Alternatively, we could have used an approach utilizing whole-brain coverage with very high levels of acceleration, where a dual-polarity readout is used to mitigate the “fuzzy-ripple” artifacts that arise from gradient imperfections (Huber et al., 2025). Using this approach, our recent work achieves whole-brain VASO with 0.85 mm isotropic voxels and TR 4.88 per volume pair (Beckett et al., 2026).

This novel hardware is being further optimized for mesoscale imaging and will become more available at other research centers. Outstanding challenges include expanding spatial coverage without sacrificing resolution or sensitivity, which should allow imaging both hippocampi within a study for a fuller characterization of memory function, and functional connectivity analyses to infer information flow between close (e.g. entorhinal cortex, Koster et al., 2018; Zhang et al., 2023, or parahippocampus, Warrington et al., 2025) and distant neocortical regions involved in primary sensory and executive control networks. Further, extending this framework downward to lower subcortical networks like the thalamus or cerebellum is not a straightforward translation.

These deep regions introduce harsh physical and structural roadblocks such as rapid arterial transit times violating our TI_1_ = 939 ms tissue-nulling window, severe B_0_ inhomogeneity gradients near the skull base, and pronounced physiological motion across the interleaved 3.2 s volume pairs. The approaches used for high-resolution VASO imaging of the hippocampus have also been used for imaging of deep-brain regions such as amygdala and cerebellum (Huber et al., 2024), and the extra performance allowed by the NexGen will offer further opportunities for optimization in these regions.

### Impact of layer-fMRI with reduced venous bias in the hippocampus in future applications

Layer-resolution accuracy is central to hippocampal investigations aiming to, for example, dissociate pathways terminating in distinct deep and superficial laminae of subfield CA1, or to link these circuits to laminar-specific pathology in various neurodegenerative conditions. This specificity across cortical depths is where VASO holds a specific advantage over BOLD contrast. The macroscopic venous contamination inherent to BOLD introduces noise and complicates the interpretation of layer-specific statistics, potentially misleading neuroscientific conclusions on for example the directionality of information flow.

While algorithmic models designed to remove venous bias in BOLD exist (Fracasso et al., 2018; Kashyap et al., 2018), they have primarily been developed for the well-studied venous draining patterns of the neocortex. However, the hippocampus and associated allocortical/subcortical structures are geometrically complex and exhibit individually variable venous architecture (Duvernoy et al., 2005; Haast et al., 2024). In these regions, approaches to improve BOLD depth accuracy remain limited, for example exclusion of voxels with strongest venous signals (Pfaffenrot et al., 2025). Collection of high-resolution data to map the vasculature (e.g., Gulban et al., 2026) can also aid in the interpretation of cortical layer profiles collected using high-resolution BOLD (Pfaffenrot et al., 2025; Pizzuti et al., 2026).

Despite its spatial specificity, the use of VASO for layer-fMRI can be limited by its reduced tSNR compared to BOLD (**Supplementary Figure S3**) and lower detection sensitivity in individual-level statistics (Huber et al., 2017). This lower raw sensitivity often necessitates longer acquisitions, while the relatively long effective TRs required for interleaved blood-nulling inherently limit its temporal resolution. Together these factors can, in practice, restrict the types of experimental designs and inquiries accessible in cognitive and clinical neuroscience (e.g., fast event-related paradigms). While advanced denoising strategies such as NORDIC have emerged as promising solutions to improve VASO sensitivity, these post-processing strategies can strongly modulate layer-specific statistics near tissue boundaries (Guidi et al., 2026; Knudsen et al., 2025). Thus, highly principled approaches are required if statistics rely on aggressive denoising to uncover delicate layer-resolution effects. Given these trade-offs, a promising path forward is a dual-contrast cross-validation framework, where BOLD statistics are validated in a representative subsample of participants using non-BOLD or spin-echo variants (Muckli et al., 2015). Notably, the VASO sequence adapted here allows simultaneous collection of VASO and BOLD for such validation.

Looking beyond the hippocampus, the tools developed here provide a starting point for mapping layer-specific connections within lower brain areas of cortex and between neo-cortex and sub-cortex, in the context of cognitive neuroscience, clinical neuroscience, and neurology. Until now, more than 95% of all >300 published human layer-fMRI studies could solely focus on brain areas in the upper half of the brain (source: layerfmri.com/papers), limiting the potential to capture neural information flow across laminar microcircuits throughout the entire human brain. The methodology developed here helps the field overcome these limitations and finally address questions of many influential theories of brain function that posit laminar signals with origins and destinations in distinct cortical layers and lower brain areas, such as such as predictive coding, thalamocortical loop models, limbic–cortical integration models for emotion and decision-making.

We expect that the ever-advancing hardware and software tools and improvements of high-resolution fMRI will ultimately transform our understanding of cognition in the awake, behaving human brain. Accurate laminar imaging of the hippocampus opens the door to investigating computational mechanisms behind any number of neuropsychological phenomena. Imaging layers of the hippocampus are critical for investigating neural correlates of episodic memory (Koster et al., 2018) and spatial navigation (Ahmadi et al., 2024). Fine-grained hippocampal dysfunction is also associated with various clinical conditions including, including early damage to CA1 and entorhinal cortex layer II in Alzheimer’s disease (Gómez-Isla et al., 1996; Kerchner et al., 2012), epilepsy related to hippocampal dyslamination (Aitken et al., 2025), mood and disorders related to CA3 dendrites and DG (Leal and Yassa, 2018; Rutland et al., 2019), and hippocampal prediction errors underlying hallucinations in schizophrenia (Haarsma and Kok, 2025). Imaging the layered structure of the hippocampus can be a useful research tool to capture the development and propagation of these abnormalities in the living human brain.

## Acknowledgments

Research reported in this publication was supported by the National Institutes of Health (NIH) under award numbers U01-EB025162 (NIBIB), U24-NS129949 (NINDS), and R44-MH129278 (NIMH); and by Weill Neurohub. We thank Dr An Vu for his helpful feedback and discussion.

## Data Availability Statement

The data that support the findings of this study are openly available in OpenNeuro at https://openneuro.org, reference number ds007122.

## Ethics Statement

The study was conducted in accordance with the office for protection of human subjects, UC Berkeley IRB, and all participants signed informed consent.

## Competing Interests

Authors declare no conflicts of interest.

## Notes

https://doi.org/10.18112/openneuro.ds007122.v1.1.0

### Competing Interest Statement

The authors have declared no competing interest.

### Summary of Updates

Three subjects added; ROI-based analysis included; updated figures, reporting and discussion.

