## Supplementary Materials for "Functional imaging of hippocampal layers using VASO and BOLD on the Next Generation (NexGen) 7T scanner"

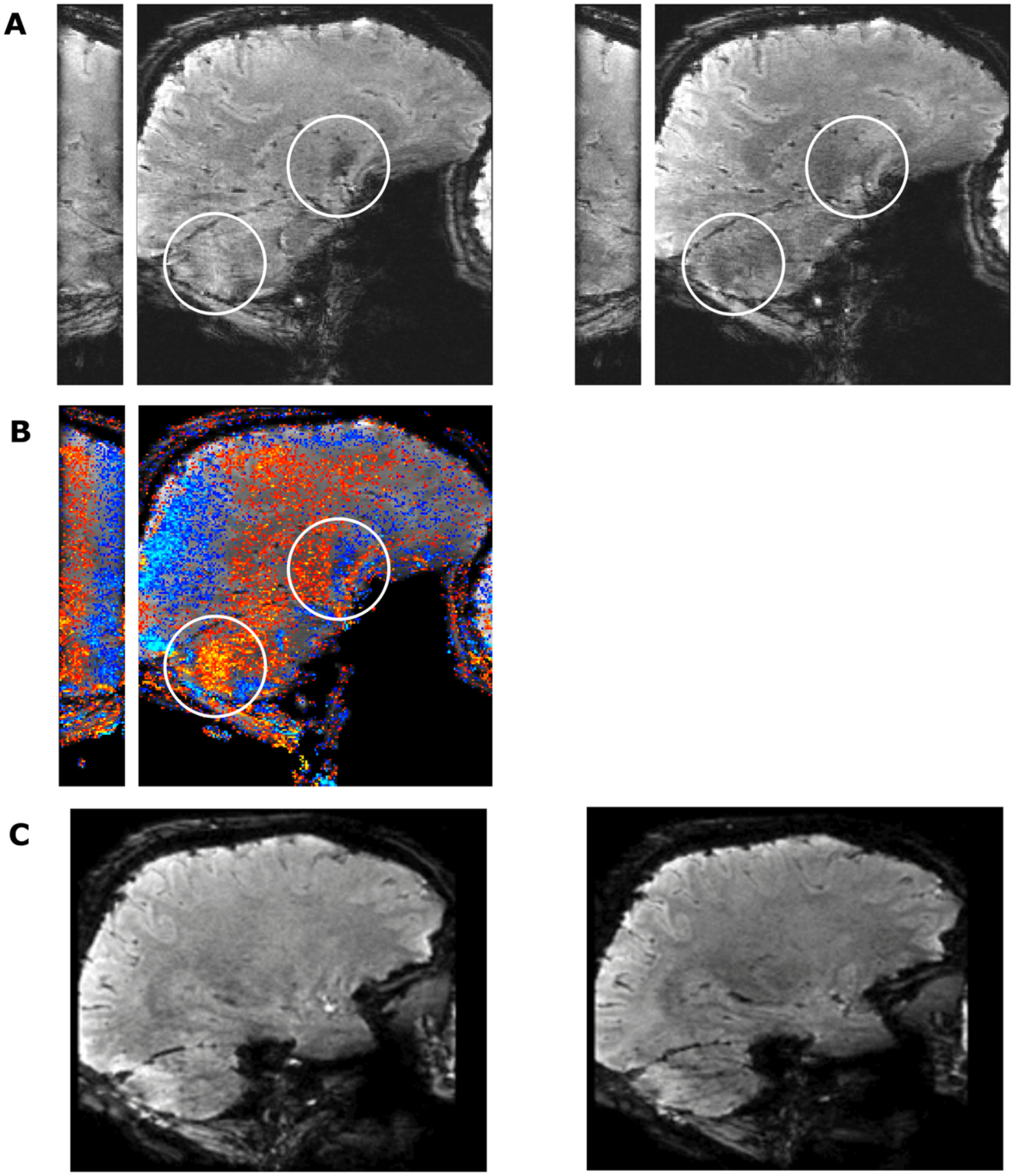


**Figure S1. Examples from initial testing stage.** (**A**) Two consecutive Nulled volumes acquired using a segmented acquisition scheme, showing a phase artifact from combining two different inversion recoveries to an image in the left-right direction. (**B**) The same phase artifact leading to artefactual activation differences in task activation z-scores. (**C**) The shorter echo spacings available on NexGen 7T scanner also helped reduce signal dropout in the medial temporal regions, as shown in the lower images.

**Table S1.** Task performance during fMRI acquisition

| Participant | AM responses | AM response time (s) | MA responses | MA response time (s) |
|---|---|---|---|---|
| S1 | 100% | 1.61 | 100% | 1.93 |
| S2 | 97.77% | 5.93 | 88.89% | 11.39 |
| S3 | NA | NA | NA | NA |
| S4 | 100% | 2.36 | 97.77% | 4.38 |
| S5 | 100% | 3.73 | 100% | 3.36 |
| S6 | 100% | 2.49 | 100% | 6.41 |
| S7 | 95.55% | 2.80 | 97.77% | 4.39 |
| S8 | 100% | 4.85 | 100% | 5.00 |
| S9 | 100% | 2.34 | 100% | 2.84 |

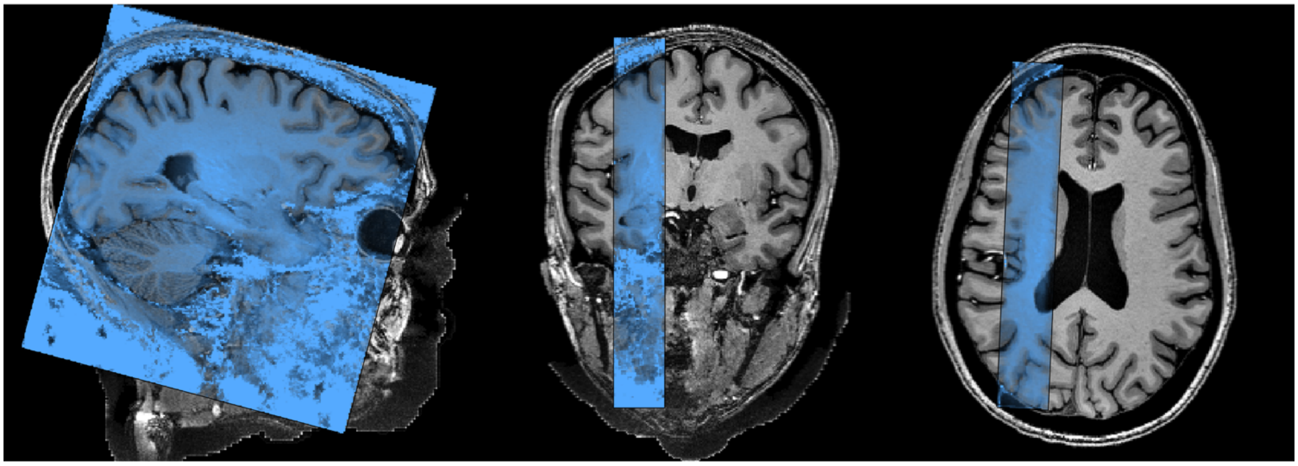

**Figure S2. Representative slab coverage and functional field of view (FOV).** Sagittal (left), coronal (middle), and axial (right) slices from a representative participant show the localized partial FOV (highlighted in blue) overlaid on the anatomical MP2RAGE.

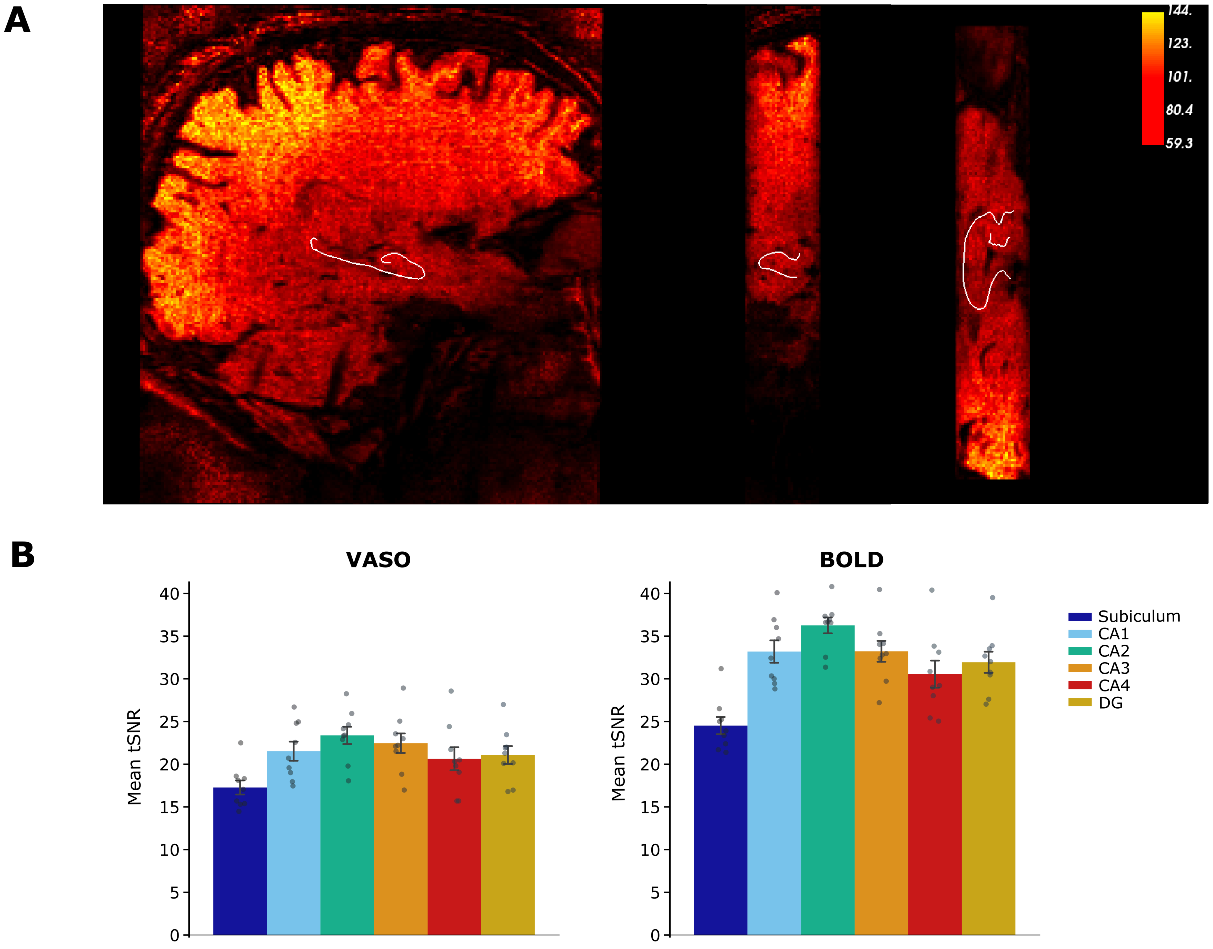


**Figure S3. Hippocampal temporal signal-to-noise ratio (tSNR) for VASO and BOLD contrasts.** (**A**) Representative sagittal, coronal, and axial tSNR maps from a single participant, extracted after motion correction and NORDIC denoising (factor error 1.0). Hippocampal outer boundary is denoted by a white line. (**B**) Mean tSNR values across subfields for VASO and BOLD (N = 9 subjects). Bar heights represent the group mean, with individual data points corresponding to subject-specific means averaged across functional runs. Error bars denote standard error of the mean (SEM) across subjects.

**Table S2. 3-Way Repeated Measures ANOVA and post-hoc pairwise comparisons for memory task activation (AM > MA).** (**A**) Omnibus 3-way repeated measures ANOVA evaluating the main effects and interactions of longitudinal Segment (Anterior, Middle, Posterior), Layer (Inner, Middle, Outer), and Modality (BOLD, VASO). (**B**) Follow-up post-hoc pairwise longitudinal segment comparisons. (**C**) Follow-up post-hoc pairwise laminar layer comparisons. All tests represent a repeated measures design across a sample size of $N = 9$ subjects. Multiple comparisons for post-hoc tests were corrected using the Holm-Bonferroni method.

| **Part A: Omnibus 3-Way Repeated Measures ANOVA** | | | | | | | |
|---|---|---|---|---|---|---|---|
| Effect | F | ddof1 | ddof2 | p | | | |
| Segment | 4.453 | 2 | 16 | 0.029 | * | | |
| Layer | 9.058 | 2 | 16 | 0.002 | * | | |
| Modality | 11.466 | 1 | 8 | 0.010 | * | | |
| Segment × Layer | 0.561 | 4 | 32 | 0.692 | | | |
| Segment × Modality | 4.414 | 2 | 16 | 0.030 | * | | |
| Layer × Modality | 7.304 | 2 | 16 | 0.006 | ** | | |
| Segment × Layer × Modality | 0.185 | 4 | 32 | 0.945 | | | |
| **Part B: Post-Hoc Pairwise Longitudinal Segment Comparisons** | | | | | | | |
| Modality | A | B | t | dof | p | $p_{Holm}$ | |
| BOLD | Anterior | Middle | 1.520 | 8 | 0.167 | 0.167 | |
| BOLD | Middle | Posterior | 2.008 | 8 | 0.080 | 0.159 | |
| BOLD | Anterior | Posterior | 3.191 | 8 | 0.013 | 0.038 | * |
| VASO | Anterior | Middle | 0.635 | 8 | 0.543 | 1.000 | |
| VASO | Middle | Posterior | 0.488 | 8 | 0.639 | 1.000 | |
| VASO | Anterior | Posterior | 1.494 | 8 | 0.174 | 0.521 | |
| **Part C: Post-Hoc Pairwise Layer Comparisons** | | | | | | | |
| Modality | A | B | t | dof | p | $p_{Holm}$ | |
| BOLD | Inner | Middle | 3.263 | 8 | 0.011 | 0.034 | * |
| BOLD | Middle | Outer | 2.877 | 8 | 0.021 | 0.034 | * |
| BOLD | Inner | Outer | 3.145 | 8 | 0.014 | 0.034 | * |
| VASO | Inner | Middle | 0.836 | 8 | 0.428 | 0.428 | |
| VASO | Middle | Outer | 2.287 | 8 | 0.052 | 0.155 | |
| VASO | Inner | Outer | 1.980 | 8 | 0.083 | 0.166 | |

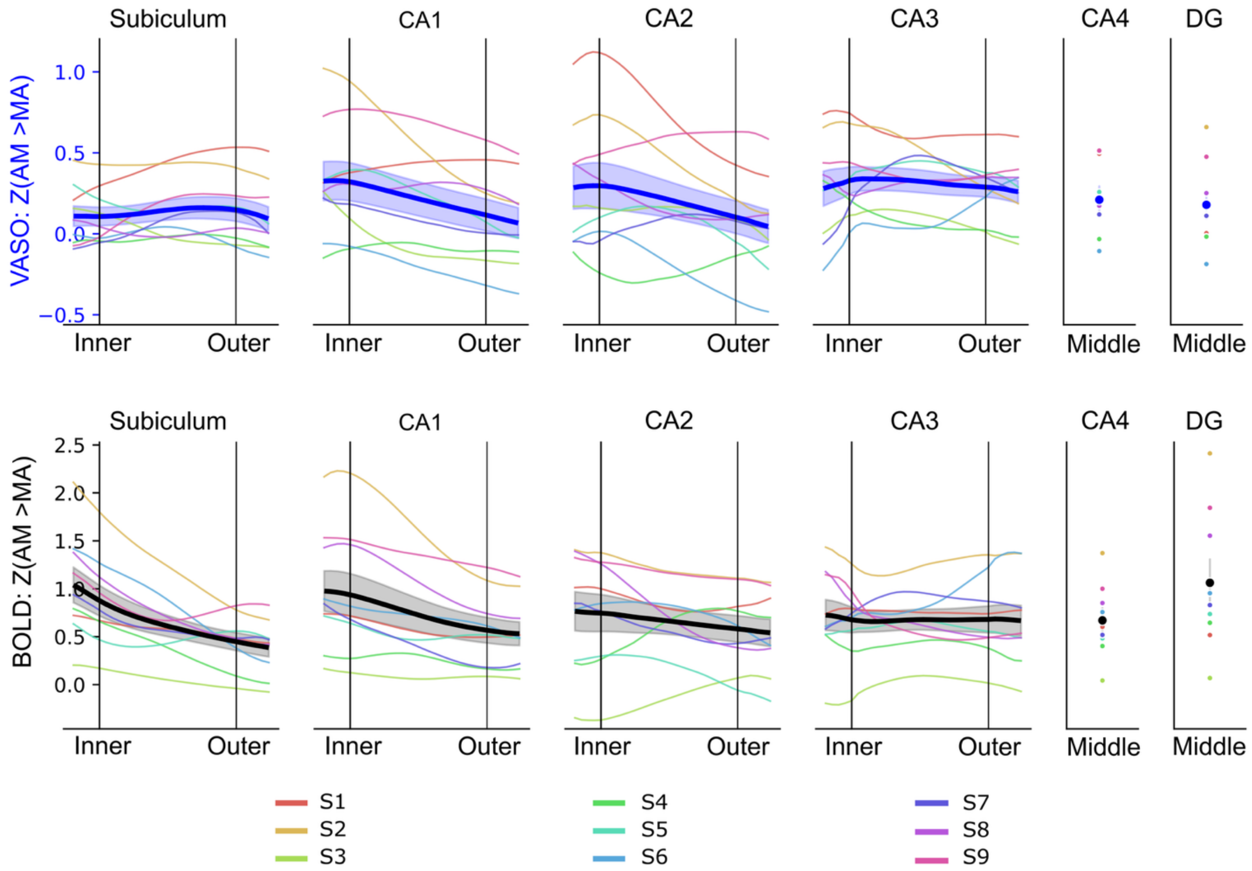


**Figure S4. Layer-specific memory task activation (AM > MA) based on VASO and BOLD.** ROI analysis of layer activation per hippocampal subfield (N = 9).

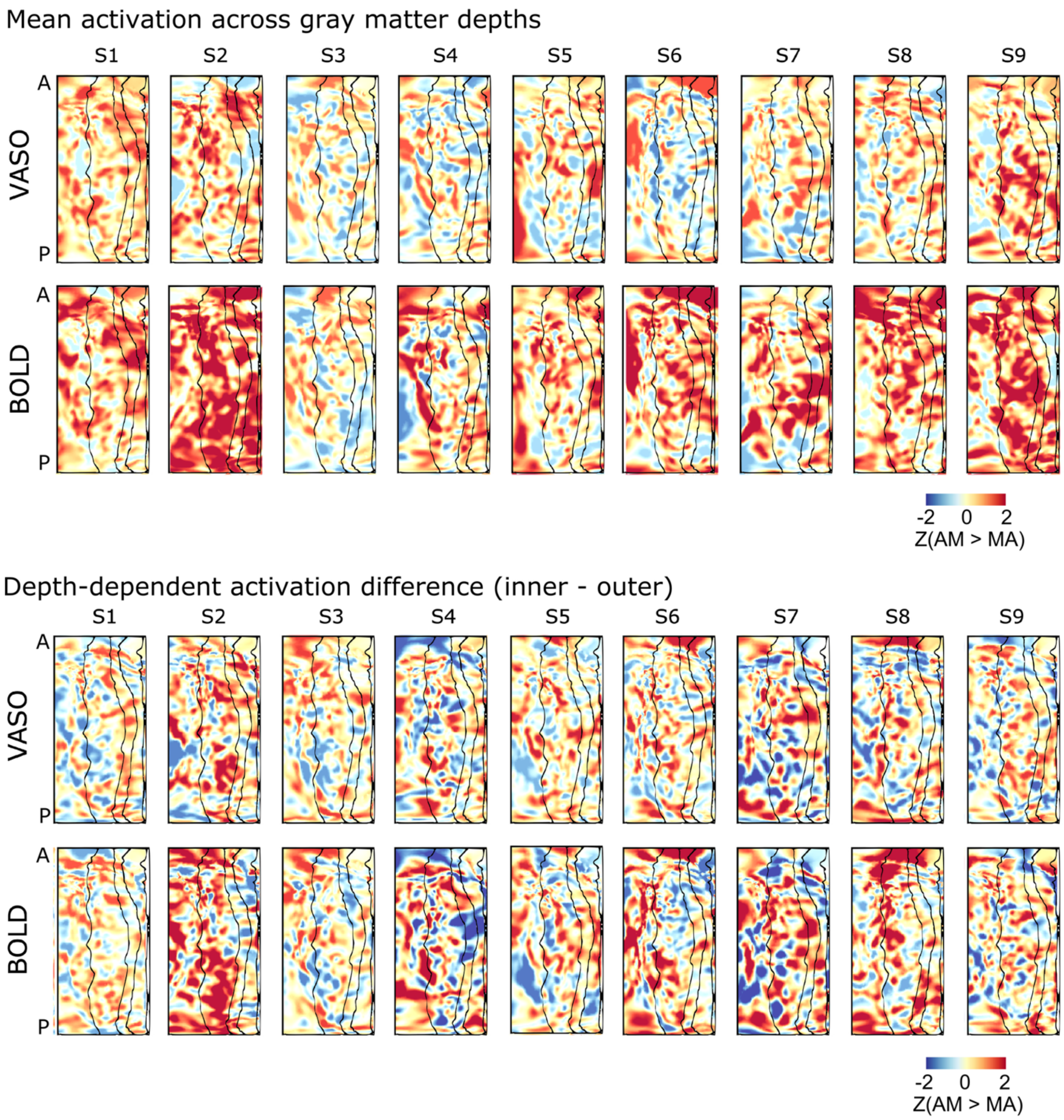


**Figure S5. Individual-level unthresholded task activation maps (AM > MA).** Top block displays mean depth profiles, and bottom block displays laminar difference profiles (inner - outer) across all participants. To preserve the high-resolution spatial topography, maps are displayed with minimal surface smoothing (1 mm FWHM).

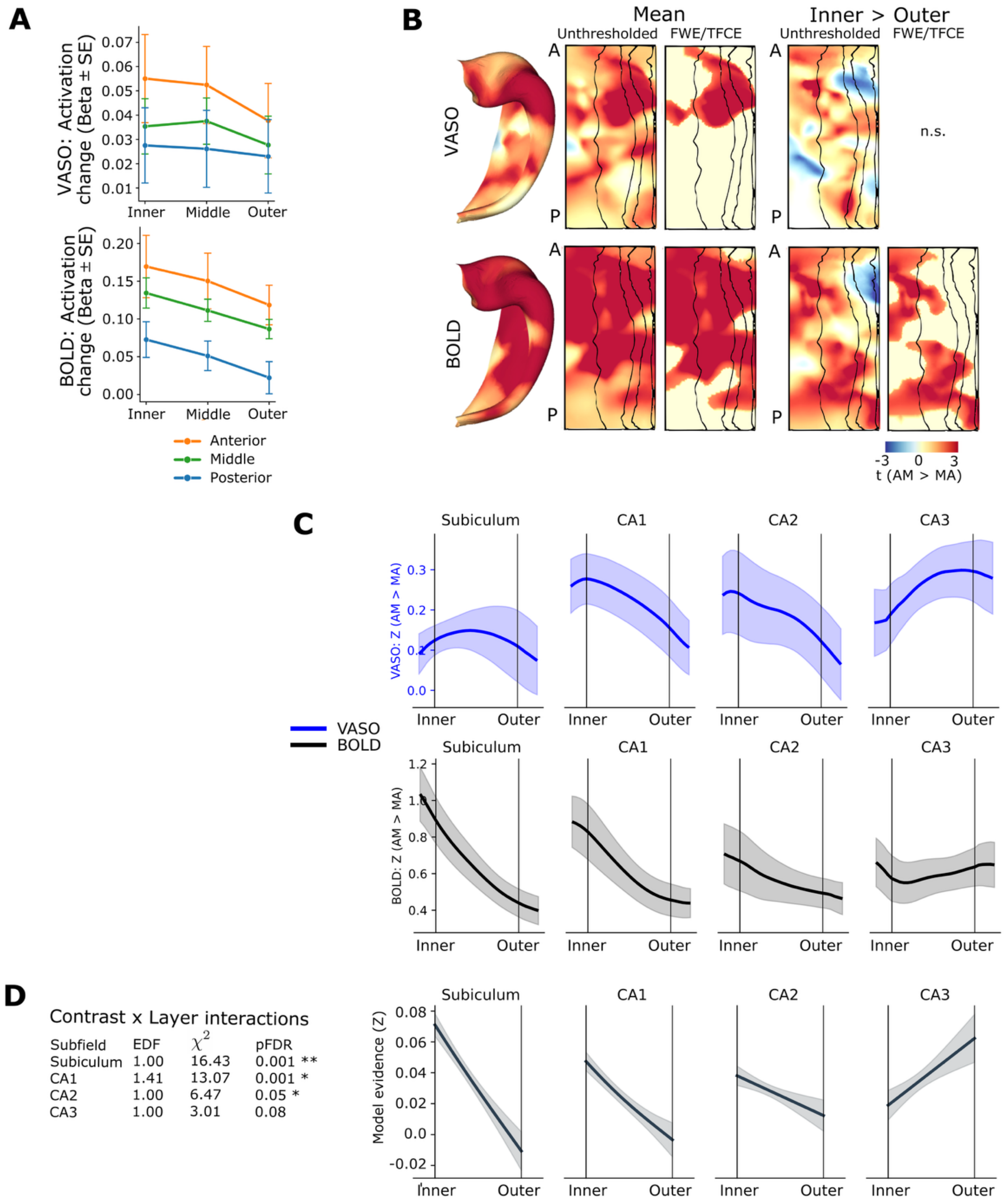


**Figure S6. Control analysis of hippocampal memory task activation (AM > MA) based on VASO and BOLD with aCompCor covariates for physiological noise.** (**A**) Memory task activation (AM > MA) tracked across longitudinal (anterior, middle, posterior) and depth (inner, middle, outer) bins. 3-way repeated measures ANOVA (Layer, Segment, Modality) confirmed significant interactions of Layer × Modality and Segment × Modality. (**B**) Group activation patterns visualized on unfolded hippocampal surfaces (5 mm surface smoothing). Within each modality, unthresholded t-statistic maps are displayed side-by-side with family-wise error rate thresholded maps. Panels display general activation topology averaged across gray matter depth alongside maps contrasting inner and outer depths. (**C**) Mean activation depth profiles (z-scores) plotted continuously for VASO (blue) and BOLD (black) across anatomical subfields. (**D**) ROI-based PTA analysis, comparing the shapes of VASO and BOLD layer profiles using multilevel smoothing splines, revealed significant differences in subiculum, CA1 and CA2. Curves show model evidence for depth-dependent contrast differences.

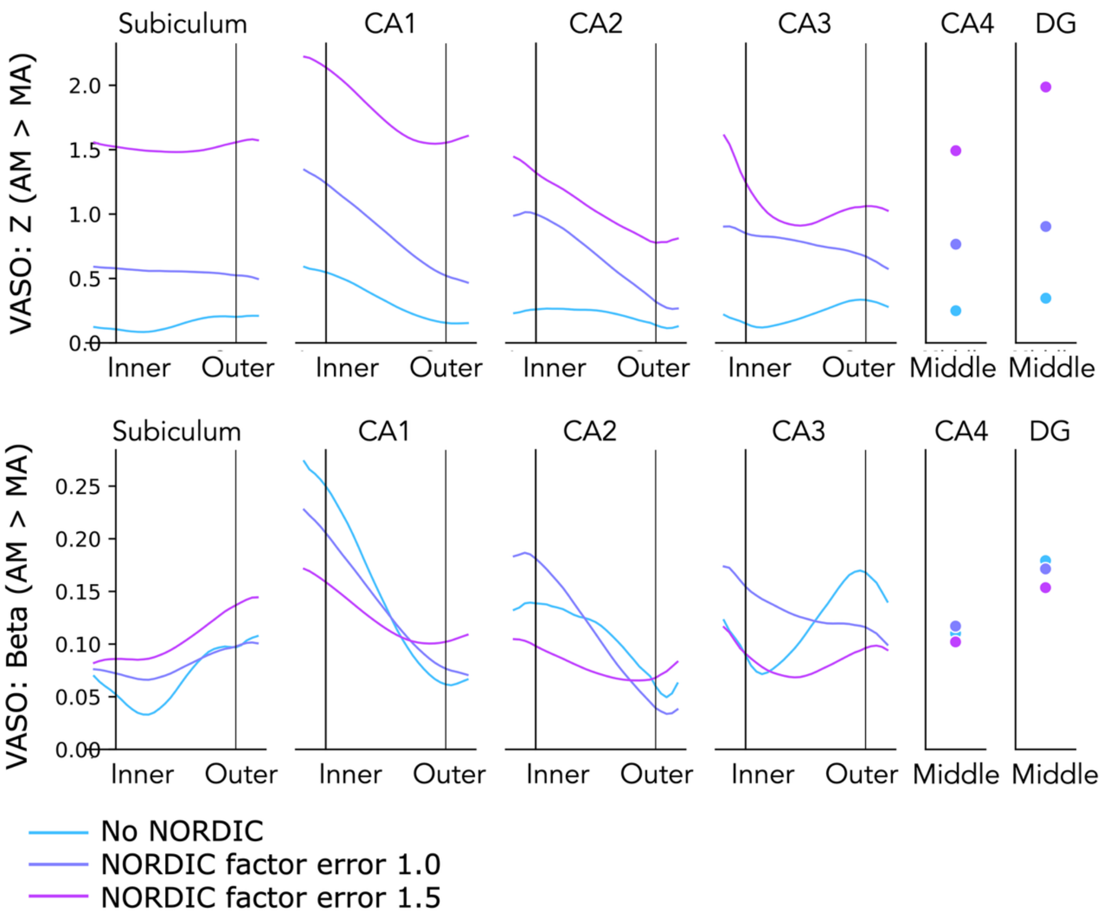


**Figure S7. VASO layer profiles (AM > MA) obtained at different factor error levels of NORDIC, in an example participant.** NORDIC increased z-score profiles without fundamentally changing the relative profile shape, but some beta estimates (e.g. inner layers of CA1) were reduced. This control analysis included no nuisance regressors.

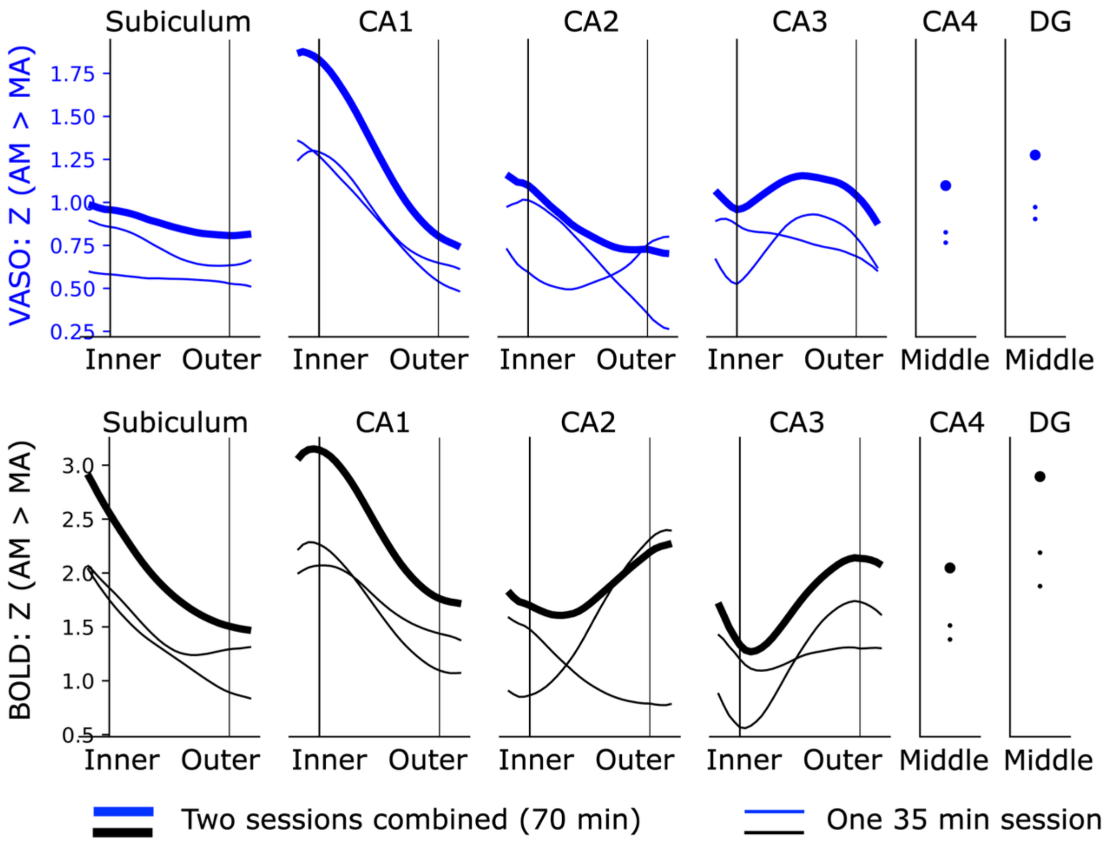


**Figure S8. Reliability of the layer profiles in repeat testing in one participant.** Layer-specific memory task activation (AM > MA) based on VASO and BOLD from the same acquisition, showing similar profile shapes in subiculum and CA1 during two sessions acquired at different days and their combined analysis. This control analysis included no nuisance regressors.

**Table S3. 3-Way Repeated Measures ANOVA and post-hoc pairwise comparisons for memory stage activation (elaboration > reconstruction).** (**A**) Omnibus 3-way repeated measures ANOVA evaluating the main effects and interactions of longitudinal Segment (Anterior, Middle, Posterior), Layer (Inner, Middle, Outer), and Modality (BOLD, VASO). (**B**) Follow-up post-hoc pairwise longitudinal segment comparisons. (**C**) Follow-up post-hoc pairwise laminar layer comparisons. All tests represent a repeated measures design across a sample size of $N = 8$ subjects. Multiple comparisons for post-hoc tests were corrected using the Holm-Bonferroni method.

| **Part A: Omnibus 3-Way Repeated Measures ANOVA** | | | | | | | |
|---|---|---|---|---|---|---|---|
| Effect | F | ddof1 | ddof2 | p | | | |
| Segment | 7.236 | 2 | 14 | 0.007 | ** | | |
| Layer | 1.276 | 2 | 14 | 0.310 | | | |
| Modality | 0.362 | 1 | 7 | 0.567 | | | |
| Segment × Layer | 1.777 | 4 | 28 | 0.162 | | | |
| Segment × Modality | 0.880 | 2 | 14 | 0.437 | | | |
| Layer × Modality | 0.888 | 2 | 14 | 0.433 | | | |
| Segment × Layer × Modality | 1.674 | 40 | 28 | 0.184 | | | |
| **Part B: Post-Hoc Pairwise Longitudinal Segment Comparisons** | | | | | | | |
| Modality | A | B | t | dof | p | $p_{Holm}$ | |
| BOLD | Anterior | Middle | 0.003 | 7 | 0.998 | 0.998 | |
| BOLD | Middle | Posterior | -2.234 | 7 | 0.061 | 0.121 | |
| BOLD | Anterior | Posterior | -4.080 | 7 | 0.005 | 0.014 | |
| VASO | Anterior | Middle | -0.032 | 7 | 0.976 | 0.976 | |
| VASO | Middle | Posterior | -3.053 | 7 | 0.019 | 0.037 | * |
| VASO | Anterior | Posterior | -4.250 | 7 | 0.004 | 0.011 | * |
| **Part C: Post-Hoc Pairwise Layer Comparisons** | | | | | | | |
| Modality | A | B | t | dof | p | $p_{Holm}$ | |
| BOLD | Inner | Middle | 1.766 | 7 | 0.121 | 0.362 | |
| BOLD | Middle | Outer | 0.864 | 7 | 0.416 | 0.424 | |
| BOLD | Inner | Outer | 1.374 | 7 | 0.212 | 0.424 | |
| VASO | Inner | Middle | 0.635 | 7 | 0.545 | 1.000 | |
| VASO | Middle | Outer | 0.055 | 7 | 0.958 | 1.000 | |
| VASO | Inner | Outer | 0.185 | 7 | 0.859 | 1.000 | |

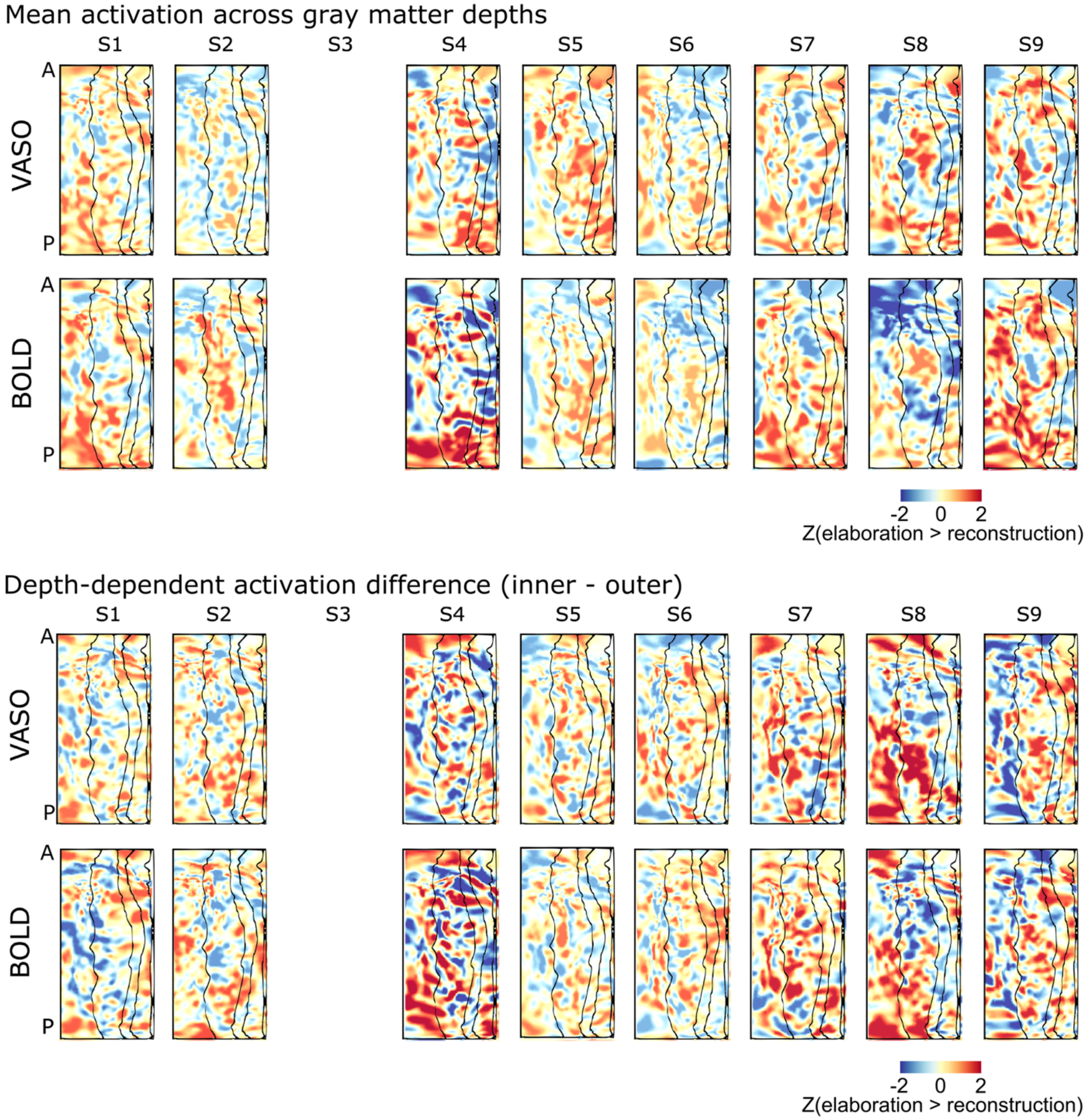


**Figure S9. Individual-level unthresholded task activation maps (elaboration > reconstruction).** Top block displays mean depth profiles, and bottom block displays laminar difference profiles (inner - outer) across all participants. To preserve the high-resolution spatial topography, maps are displayed with minimal surface smoothing (1 mm FWHM).

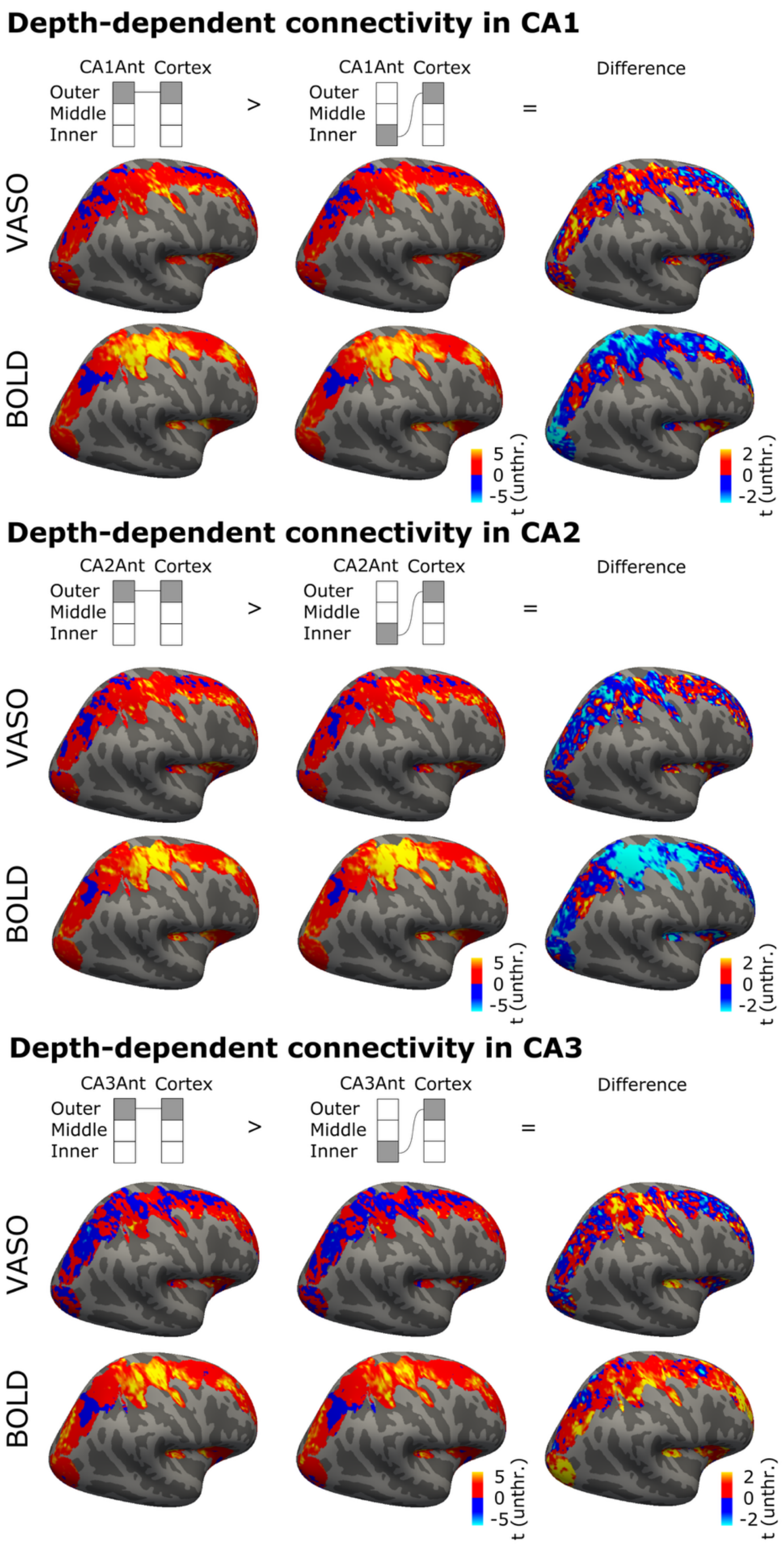


**Figure S10. Hippocampal depth-dependent functional connectivity in CA1, CA2 and CA3 based on VASO and BOLD**. Connectivity is seeded by the inner and outer depths of the anterior third along the long axis. Statistics are shown only for the right hemisphere vertices imaged in all nine participants. All statistics are shown without thresholding or correction for multiple comparisons.
